# Genome-scale modelling and stable isotope tracing reveal dynamic metabolic reprogramming in a human cardiac hypertrophy model

**DOI:** 10.64898/2026.11.06.764817

**Authors:** Catherine Keiko Takemura-James, Katarzyna Kmiotek-Caller, Ravi A. Kumar, Eliano dos Santos, Rafael R. Oexner, Christina M. Reumiller, Sharwari Verma, Ella Bauwens, Elisavet Kiaitsi, Celio X. C. Santos, Katrin Streckfuss-Bömeke, Ajay M. Shah, Anna Zoccarato

## Abstract

Pathological cardiac hypertrophy is associated with metabolic remodelling; however the temporal dynamics of this remodelling remain poorly defined in human cardiomyocyte systems. Here, we combined transcriptomic profiling, genome-scale metabolic modelling and stable isotope-resolved metabolomics (SIRM) to map the metabolic consequences of endothelin-1 (ET-1) stimulation in human iPSC-derived cardiomyocytes (hiPSC-CMs). Integration of RNA sequencing data obtained after 8 and 48 hours of ET-1 treatment with the Human2.0 genome-scale metabolic model predicted time-dependent remodelling, characterised by progressive increases in glycolytic flux, enhanced glucose carbon entry into the TCA cycle, and a decrease in fatty acid β-oxidation. Computational flux predictions were independently confirmed by extracellular flux analysis and ¹³C-substrate tracing, with concordance between modelled and measured fluxes. Extracellular flux analysis showed increased glycolytic capacity and mitochondrial respiration after 48 hours, with greater sensitivity to inhibition of mitochondrial pyruvate transport. Glucose tracing revealed increased ¹³C enrichment of glycolytic, pentose phosphate pathway, serine-glycine and TCA cycle metabolites, consistent with altered glucose-derived carbon allocation. Increased m+3 labelling of selected TCA-related metabolites was compatible with pyruvate anaplerosis, although pathway-specific fluxes were not resolved. In three-dimensional engineered heart tissue (EHTs) treated with ET-1 for 7 days, increased hypertrophic marker expression was accompanied by a blunted inotropic response, and metabolomics showed similar changes in metabolite abundance and glucose-derived labelling to 2D cultures. These findings support hiPSC-CMs and EHTs as complementary platforms for studying early metabolic remodelling in pathological cardiac hypertrophy and provide an integrated framework linking model predictions to functional and isotope-tracing measurements.

**Highlights:**

- Genome-scale modelling of RNA sequencing from ET-1-stimulated human iPSC-derived cardiomyocytes at 8 and 48 hours predicts progressive glycolytic activation, increased glucose carbon entry into the TCA cycle, and decreased fatty acid β-oxidation.
- Extracellular flux analysis and ¹³C-substrate tracing confirm the modelled fluxes with increased partitioning into the pentose phosphate, aspartate and glutamate synthesis, and serine-glycine pathways, suggesting anabolic glucose use.
- Metabolomics in ET-1 treated engineered heart tissues recapitulates the iPSC-CM signature, mirroring maladaptive remodelling and establishing both systems as platforms for resolving early hypertrophic flux changes.

## 1.0 Introduction

Pathological cardiac hypertrophy is characterised by re-activation of the foetal gene programme, alterations to intracellular signalling, and a substantial metabolic remodelling. Unlike physiological hypertrophy, the initial compensatory response can become maladaptive and contribute to heart failure. A hallmark of this metabolic remodelling is a substrate switch away from the fatty acid-dependent oxidative metabolism of the healthy heart, towards increased glucose utilisation and reduced metabolic flexibility[1–3]. Importantly, the pattern of glucose utilisation appears to evolve with disease progression.

Glucose uptake and glycolysis have been shown to be increased in pathological hypertrophy[4–6], often with enhanced glucose oxidation during the compensated phase[5, 7], whereas in established heart failure, glycolysis may become uncoupled from mitochondrial glucose oxidation, with accumulation of glycolytic intermediates and impaired energy provision[8–12]. It is increasingly appreciated that increased glucose uptake in hypertrophy not only contributes to higher energetic needs but may also serve biosynthetic requirements by directing carbon flux into pathways that supply nucleotides, amino acids, and one-carbon units for cardiomyocyte remodelling[13–15]. Human studies have identified substantial metabolic alteration in failing myocardium[11, 16]. However, changes during early hypertrophic remodelling remain incompletely characterised in human cardiomyocytes. Whilst animal models have established important principles[5, 9, 10, 12, 17], species differences in substrate preference, metabolic rate, and mitochondrial density could influence cellular metabolism[18–20], motivating complementary studies in human systems[21, 22].

Human iPSC-derived cardiomyocytes (hiPSC-CMs) have emerged as a powerful human-relevant model that recapitulates key features of pathological hypertrophy including myocyte disarray, foetal gene re-expression, and natriuretic peptide secretion in response to neurohumoral stimulation[23–26]. We have previously demonstrated that prolonged culture of hiPSC-CMs results in metabolic maturation[27], providing a more physiologically relevant baseline from which to study disease-induced remodelling. Nevertheless, the metabolic flux changes accompanying hypertrophic remodelling in hiPSC-CMs have not been characterised.

Genome-scale metabolic models (GEMs) offer a stoichiometric framework for integrating transcriptomic data to predict network-wide flux distribution[28–30], while stable isotope-resolved metabolomics (SIRM) enables direct experimental tracing of carbon partitioning through metabolic pathways. The combination of these approaches has recently been applied to human non-ischaemic cardiomyopathy, using an animal model for validation with ^13^C tracer studies[31]. However, whether this integrated approach can resolve the flux-level changes occurring during the earlier hypertrophic remodelling phase, and whether it is directly applicable in human iPSC-CM and engineered heart tissue (EHT) platforms, remains unexplored. Here, we employed an integrated approach combining RNA sequencing, genome-scale metabolic modelling using Human2.0, extracellular flux analysis, and SIRM to examine ET-1-induced metabolic reprogramming in hiPSC-CMs. We assess time-dependent model predictions and compare them with respiratory measurements and ^13^C-glucose and ^13^C-palmitate labelling after 48 hours of ET-1 treatment. We also examined metabolic and contractile changes in EHTs after 7 days of ET-1 treatment. By linking genome-scale model predictions with experimental SIRM data, this study provides a comprehensive, human-relevant characterisation of ET-1-induced early metabolic reprogramming and establishes an integrated framework for investigating metabolic remodelling in hiPSC-CM models.

## 2.0 Materials and Methods

### 2.1 Cell culture and hiPSC differentiation

The hiPSC lines used in the study, CRTL2 and WT4, were derived from healthy donors and were described and characterized previously[32]. hiPSCs were maintained in mTesR Plus medium (StemCell Technologies; Vancouver, Canada) on hESC-Qualified Matrigel®-coated flasks (Corning Life Sciences, Tewksbury, USA) with daily medium changes. Once cultures reached 70-80% confluence, cells were passaged using 0.5 mM EDTA solution (Invitrogen; Thermo Fisher Scientific; Waltham, USA).

Human foreskin fibroblasts (HFF-1) were purchased from ATCC (Manassas, USA) and cultured in DMEM high glucose (Sigma-Aldrich; Merck Life Science; Burlington, USA), supplemented with 15% foetal bovine serum (FBS; Sigma-Aldrich). At 90-95% confluence, cells were split using TrypLE Express (Gibco; Thermo Fisher Scientific). Cell cultures were maintained at 37 °C in a humidified incubator with 5% CO₂.

hiPSCs were differentiated into cardiomyocytes (hiPSC-CMs) using a protocol of Wnt modulation and metabolic selection as described previously [33, 34](Supplementary Figure 1A) and maintained in a cardiac culture medium[33]. Differentiation efficiency and quality of hiPSC-CMs was assessed by flow cytometry. Cells were stained for the cardiomyocyte marker cardiac troponin T (cTnT; Abcam, Cambridge, UK; AntiCardiac Troponin T [1C11], ab8295) and the fibroblast marker TE7 (Sigma-Aldrich; AntiFibroblasts Antibody, clone TE7, CBL271) (Supplementary Figure 1B). Only cultures with ≥85% cTnT-positive cells and ≤5% TE7-positive cells were included in subsequent experiments. To evaluate the effect of endothelin-1 (ET-1; Sigma-Aldrich) over time, hiPSC-CMs were incubated with 10 nM ET-1 for 8 or 48 hours and compared to untreated controls.

### 2.2 Cell volume measurement via flow cytometry

Cell size was determined by flow cytometry as previously described[35]. In brief, untreated or ET-1-treated hiPSC-CM were dissociated with TrypLE Express Enzyme (Thermo Fisher Scientific) and resuspended in PBS containing 0.5% bovine serum albumin (BSA) and held on ice until acquisition. Samples were run on a BD FACSymphony A1 cytometer (BD Biosciences, Franklin Lakes, NJ, USA), and forward scatter area (FSC-A) was converted to cell diameter using a standard curve constructed from beads of defined size (Flow Cytometry Size Calibration Kit, Thermo Fisher Scientific) (Supplementary Figure 1C). Up to 50,000 events were recorded per condition in BD FACSDiva v9.0 (BD Biosciences), and the resulting data were analysed in FlowJo v10.0 (BD Biosciences).

### 2.3 Immunofluorescence and cell size measurement

hiPSC-CMs were seeded at 3 × 10⁴ cells per well on µ-Slide 8-well ibiTreat chamber slides (ibidi, Gräfelfing, Germany), allowed to recover for one week, then treated with ET-1 for 48 h or left untreated. Cells were fixed in 4% PFA (15 min, RT), permeabilised in 0.1% Triton X-100/PBS (15 min), blocked for 30 min in PBS with 10% goat serum and 1% BSA and stained with anti-α-actinin (Abcam, ab68194) and DAPI Merck Life Science UK Limited). Single focal planes were acquired on an Andor BC43 spinning-disc confocal microscope (Oxford Instruments, High Wycombe, UK) with a Nikon CFI Plan Apochromat Lambda D 10×/0.45 NA air objective (Nikon Instruments, Tokyo, Japan). Images were analysed in Fiji (ImageJ).

### 2.4 Engineered heart tissues (EHTs) generation and measurements of contractile force

EHTs were generated as previously described [36]. The protocol was adapted for use with the CUORE system (Optics11 Life; Amsterdam, Netherlands), a pillar-based system which integrates fibre-optic sensing and electrical pulse stimulation to enable real-time measurement of contractile activity[37]. Briefly, hiPSC-CMs and HFF-1 cell suspensions were obtained by enzymatic dissociation using Accutase (Thermo Fisher Scientific) and TrypLE Express (Thermo Fisher Scientific), respectively. For each EHT, a hydrogel mixture was prepared by combining acid-soluble TeloCol6 bovine collagen (6 mg/mL; Advanced Biomatrix, Carlsbad, USA) with an equal volume of 2× RPMI medium (Thermo Fisher Scientific), followed by addition of Neutralization Solution (Advanced Biomatrix). A 70:30 hiPSC-CM:HFF-1 cell suspension reconstituted in EHMM+T medium [36] was incorporated into the hydrogel, and the resulting mixture was dispensed into CUORE moulds pre-coated with Pluronic F127 (Sigma Aldrich). EHT constructs were incubated at 37 °C in a humidified incubator with 5% CO₂ for 1 h to allow polymerisation, after which each EHT was supplemented with 1.5 mL of EHMM+T medium. Medium was changed daily. To induce hypertrophy, EHTs were treated with 10 nM ET-1 for 7 days. After 7 days incubation, EHTs were subjected to acute β-adrenergic stimulation with isoproterenol (1 nM and 10 nM) under electrical stimulation. Contractile peaks and parameters were analysed using CUORE Viewer software (Optics11 Life).

### 2.5 cDNA synthesis and real-time quantitative polymerase chain reaction

Total RNA was isolated from hiPSC-CM or EHT samples using the ReliaPrep™ Tissue System kit (Promega; Madison, USA) according to the manufacturer’s instructions. cDNA was synthesized using the LunaScript® RT SuperMix Kit (New England Biolabs; Ipswich, USA).

Quantitative real-time PCR (qPCR) was performed with qPCR BIO SyGreen Mix Hi-ROX (PCR Biosystems; London, UK) on a StepOnePlus™ Real-Time PCR System (Applied Biosystems; Thermo Fisher Scientific) and the comparative Ct method was used, with β2–microglobulin (B2M) levels used for normalization. Primer sequences are the following: B2M FWD: TGCTGTCTCCATGTTTGATGTATCT, B2M REV: TCTCTGCTCCCCACCTCTAAGT; ANP FWD: CAGGATGGACAGGATTGGAG, ANP REV: TCCTCCCTGGCTGTTATCTTC; BNP FWD: TTGGAAACGTCCGGGTTAC, BNP REV: GGACTTCCAGACACCTGTGG; Acta1 FWD: AGGTCATCACCATCGGCAACGA Acta1 REV: GCTGTTGTAGGTGGTCTCGTGA; MYH6 FWD: CTCCTACGCAACTGCCGATA, MYH6 REV: GACACCGTCTGGAGGGATGA; Col1a1 FWD: GATTCCCTGGACCTAAAGGTGC, Col1a1 REV: AGCCTCTCCATCTTTGCCAGCA.

### 2.6 Bulk RNA-seq Analysis

hiPSC-CMs untreated or treated with 10nM ET-1 for 8 or 48 hours (n = 4, 4, 3 respectively) were detached, pelleted, and stored at −80°C. These cell pellets were transported to GENEWIZ for further processing, including RNA extraction, quality control, sequencing, and processing of raw reads. Media was aspirated, quenched in liquid nitrogen, and stored for targeted metabolomics analysis. Differential gene expression analysis was performed in R (v. 4.4.2) with DESeq2[38] on ComBat-seq[39, 40] corrected counts, with untreated baseline samples as controls. A regularised-log transformation (rlog) was applied for principal component analysis (PCA) with prcomp and visualisation. Gene set enrichment analysis (GSEA) was performed on lists of genes ranked by the Wald statistic using the clusterProfiler[41, 42] package to identify enriched KEGG pathways. Pathways were considered significant with an absolute normalised enrichment score (NES) > 1.5 and a Benjamini-Hochberg adjusted p-value < 0.05. Over-representation analysis was performed on clustered significantly (adjusted p value < 0.05) time-dependent genes as determined by the results of a likelihood ratio test (LRT) and using the degPatterns function in the package DEGReport[43], using GO terms and Reactome pathways, where terms were considered significant with a false discovery rate < 0.05. Heatmaps were created using pheatmap[43].

### 2.7 In silico modelling of metabolism in iPSC-CMs

Constraint-based modelling was performed to predict the metabolic flux distributions of iPSC-CMs treated with ET-1. Sample-specific metabolic networks were created from a base genome-scale metabolic model, Human2.0[44]. Exchange reactions were constrained by extracellular metabolite availability as determined by their presence in the media (Supplementary Table 1) based on information provided by the manufacturer. The directionality of exchange reactions was constrained by experimentally measured uptake and secretion, as determined by targeted metabolomics of sample-specific cell culture media before and after culture. The feasibility of the resulting media-constrained models was confirmed through setting maximum and minimum flux through the non-replicative biomass maintenance reaction (MAR09931) to 10 g/gDW/h and 1 g/gDW/h respectively and inspecting the solution.

Gene expression (TPM-normalised) was integrated into the sample-specific metabolic networks to further constrain the models using the Gene Inactivity Moderated by Metabolism and Expression (GIMME) algorithm[45] as implemented in the COnstraint-Based Reconstruction and Analysis (COBRA, v. 3.0)[28] toolbox in MATLAB using the COBRA solver. Reactions were determined to be active if the corresponding gene expression was within the top quartile. To simulate maintenance of proteins, lipids, and RNA the biomass maintenance reaction excluding replication precursors was set to 50% of maximum, whilst the Required Metabolic Function (RMF) was set to maximising flux through the ATP hydrolysis reaction. Flux balance analysis (FBA) was performed to predict the distribution of fluxes through the constrained metabolic network, optimising for maximum flux through the ATP hydrolysis reaction. A schematic metabolic map of central carbon metabolism was created in Cytoscape (v. 3.10.4), where nodes represent individual metabolites and edges represent reactions. Square-root-transformed group mean reaction fluxes were integrated with the map, dictating edge thickness. Pathway activity scores were calculated as the mean absolute flux of all reactions in a pathway, as previously reported[46] and displayed as z-scores in heatmaps created with pheatmap[43].

All constraint-based modelling was performed in a high-performance computing environment; King’s Computational Research, Engineering and Technology Environment (CREATE)[47].

### 2.8 Extracellular flux analysis

Extracellular flux analysis experiments were performed using a Seahorse XF^e^24 analyser as previously described[27]. Briefly, one week before the experiment hiPSC-CMs were plated on Matrigel-coated 24-well microplates at a cell density of 5 × 10^4^ cells/well. 48 hours prior to the experiment, hiPSC-CMs were incubated with 10 nM ET-1 or left untreated. Unbuffered DMEM (D5030, Sigma) was used as the base medium for all the assays and pH was adjusted to 7.35 at 37°C (assay medium). For the glycolysis stress test, on the day of the experiment cardiac culture medium was changed to D5030 1 h before assay and the extracellular acidification rate (ECAR) was determined at baseline and in response to 10 mM Glucose followed by 2.5 µM oligomycin (Sigma, 75351), and 50 mM 2-deoxyglucose, according to the manufacturer’s protocol. For the mitochondrial stress test, the assay medium was supplemented with 10 mM glucose, 1 mM sodium pyruvate and 2 mM L-glutamine, and the oxygen consumption rate (OCR) was measured at baseline and in response to 2.5 µM oligomycin (Sigma, 75351), followed by 2 µM Carbonyl cyanide 4-(trifluoromethoxy)phenylhydrazone (FCCP) (Sigma, C2920), and 2.5 µM antimycin A (Sigma, A8674) and 2.5 µM rotenone (Sigma, R8875), according to the manufacturer’s protocols.

For the substrate oxidation stress test, the assay medium was supplemented with 10 mM glucose, 2 mM glutamine, 0.2 mM carnitine, 0.2 mM BSA-Palmitate, and OCR was measured at baseline and after injection of media or 7.5 µM UK5099 (Sigma, PZ0160); or 10 µM Bis-2-(5-phenylacetamido-1,3,4-thiadiazol-2-yl)ethyl sulfide (BPTES) (Sigma, SML0601); or 15 µM Etomoxir (Sigma, E1905), followed by oligomycin, FCCP and Antimycin A and Rotenone.

Following assay completion, cells were fixed with 4% PFA for 15 min, stained with DRAQ5 (Thermo Scientific), and imaged on an Odyssey Li-Cor Imaging System (LI-COR Biosciences, Lincoln, NE) for cell number normalisation.

### 2.9 Metabolite extraction from hiPSC-CM and EHT samples for liquid chromatography–mass spectrometry

hiPSC-CM were seeded at 6 × 10^5^ in 6-well plates. Cells were incubated with glucose-free CCM supplemented with 10 mM [U-^13^C] glucose (Cambridge Isotope Laboratories, Inc.) To optimise the duration of the [U-^13^C] glucose labelling period in iPSC-CMs a time course experiment (0 h to 48 h) was performed to determine the timing for dynamic vs. steady-state label incorporation in specific metabolic pathways. The glycolytic intermediates reached isotopic steady state rapidly, whereas TCA cycle intermediates continued to accumulate label beyond 48 hours (Supplementary Figure 2). Sixteen hours was selected as a compromise between achieving steady-state labelling in the glycolytic pools and obtaining sufficient ¹³C incorporation into TCA cycle intermediates for reliable isotopologue quantification.

After the incubation period, hiPSC-CMs were washed with ice-cold phosphate-buffered saline (PBS) and quenched with 1:1:1 chloroform:methanol:water (v/v/v). Cells were scraped off the plate, and the cell suspension was placed in a tube shaker for 20 min at 4°C, followed by centrifugation at 16 000 g at 4°C for phase separation. EHT samples were incubated with 10 mM [U-^13^C] glucose for 16 h, and snap-frozen before extraction. Metabolites were first extracted using chloroform:methanol (2:1, v/v) and subjected to three sonication cycles (8 min on / 12 min off). Samples were centrifuged at 16,000 g for 10 min at 4 °C, and the supernatant was retained. The remaining pellets were re-extracted with methanol:water (2:1, v/v), followed by a second round of sonication and centrifugation.

For all conditions the polar phases were collected and dried using a SpeedVac concentrator (Savant SPD131DDA, Thermo Fisher Scientific). The dried polar fractions were stored at −80°C until MS analysis.

### 2.10 Liquid chromatography-mass-spectrometry analysis

LC–MS analysis was performed as previously described[15]. Briefly, dried polar metabolite fractions were reconstituted in acetonitrile/water (3:2, v/v), vortexed and centrifuged at 16,000 g for 3 min before analysis on a 1290 Infinity II ultrahigh-performance liquid chromatography (HPLC) system coupled to a 6546-quadrupole time-of-flight mass spectrometer (Agilent Technologies, Santa Clara, CA). Samples were separated on a Poroshell 120 HILIC-Z column (100 × 2.1 mm, 2.7 μm; Agilent Technologies) attached to a guard column (5 × 2.1 mm, 2.7 µm) and analysed in negative ionization mode using water with 10 mmol/L ammonium acetate (Solvent A) and acetonitrile with 10 mmol/L ammonium acetate (Solvent B), both containing 5 µmol/L Infinity Lab deactivator additive (Agilent Technologies). The elution gradient used was as follows: isocratic step at 95% B for 2 min, 95–65% B in 12 min, maintained at 65% B for 3 min, then returned to initial conditions over 1 min, and then the column was equilibrated at initial conditions for 8 min. The flow rate was 0.25 mL/min; the injection volume was 1 μL, and the column oven was maintained at 30°C. Feature annotation and metabolite identification were based on accurate mass and standard retention times with a tolerance of ±5 ppm and ±0.5 min, respectively, and performed with MassHunter Profinder (version 10.0.2; Agilent Technologies) using our in-house curated metabolite library[15] based on metabolite standards (Sigma-Aldrich). Samples were analysed in a single batch with technical duplicates. Metabolite abundances in cells were normalised to the sum of the total ion chromatogram (TIC), and in EHTs to protein content (µg). Pathway enrichment analysis was performed using MetaboAnalyst 5.0[49]. ^13^C label incorporation was determined using the Batch Isotopologue Extraction workflow, corrected for natural isotope abundance, and expressed as corrected abundance percentages.

### 2.9 Statistical analysis

All data are represented as mean ± standard error of the mean (SEM). Unpaired, two-tailed Student’s *t*-test was used to compare differences between ET-1 treated vs. control groups.

For mass isotopologue distribution analysis, 2-way ANOVA followed by Šidák’s multiple comparison test. For metabolite abundance analysis statistical significance was determined using multiple unpaired t-tests, with correction for multiple comparisons using the two-stage step-up method of Benjamini, Krieger, and Yekutieli, with Q = 1%. In transcriptomics data, a Benjamini-Hochberg multiple-testing adjusted p-value of < 0.05 was considered statistically significant. Statistical analyses were performed with GraphPad Prism 9.5.1 and R for Statistical Computing (v. 4.4.2).

## 3.0 Results

### 3.1 Endothelin-1 treatment induces time-dependent transcriptomic changes in hiPSC-cardiomyocytes

ET-1 induces a distinct, time-dependent transcriptomic profile in hiPSC-CMs compared to untreated control. Multivariate analysis of the transcriptomic profiles of hiPSC-CMs untreated or treated with ET-1 for 8 or 48 hours (Figure 1A) identified distinct transcriptional profiles between the three conditions, with hour 8 showing the most profound changes (Figure 1B-C), in line with previous time-course data[25]. To understand how these different transcriptomic profiles may translate into different biological functions over time, we used a hierarchical clustering approach[50] to cluster the significantly time-dependent genes (adjusted p-value <0.05, LRT) according to their gene expression profile over time. This approach identified four clusters of gene expression patterns (Figure 1D). An over-representation analysis (ORA) was performed to identify enriched pathways for each cluster using gene ontology (GO) terms and Reactome pathways. Our results showed that the transcriptomic adaptation to ET-1 induced stress appears to begin early by 8 hours, with a large number of upregulated genes which then fall again to near pre-treatment expression levels at hour 48 (Figure 1D, cluster 1). These genes correlate to an enrichment in terms associated with intra- and inter-cellular signalling and cellular component biogenesis (Suppl. Figure 3A). In cluster 2, gene expression showed an initial decrease at 8 hours followed by an increase back to baseline level at 48 hours post treatment. Gene in this cluster were associated with catabolic processes and RNA transcription. In cluster 3, where gene expression shows an upward trend over time (Figure 1D, cluster 3), enrichment in terms associated with metabolism including metabolism, small molecule metabolic process, metabolism of proteins, phosphate-containing compound metabolic process, and lipid metabolic process was identified, suggesting that ET-1 induces a progressive enrichment of transcriptomic pathways involved in metabolism (Figure 1E). Cluster 4 showed progressively decreasing expression and enrichment of terms involving transcription and negative regulation of metabolism (Figure 1D and Suppl. Fig. 3B, cluster 4) from 8 to 48 hours.

**Figure 1.**
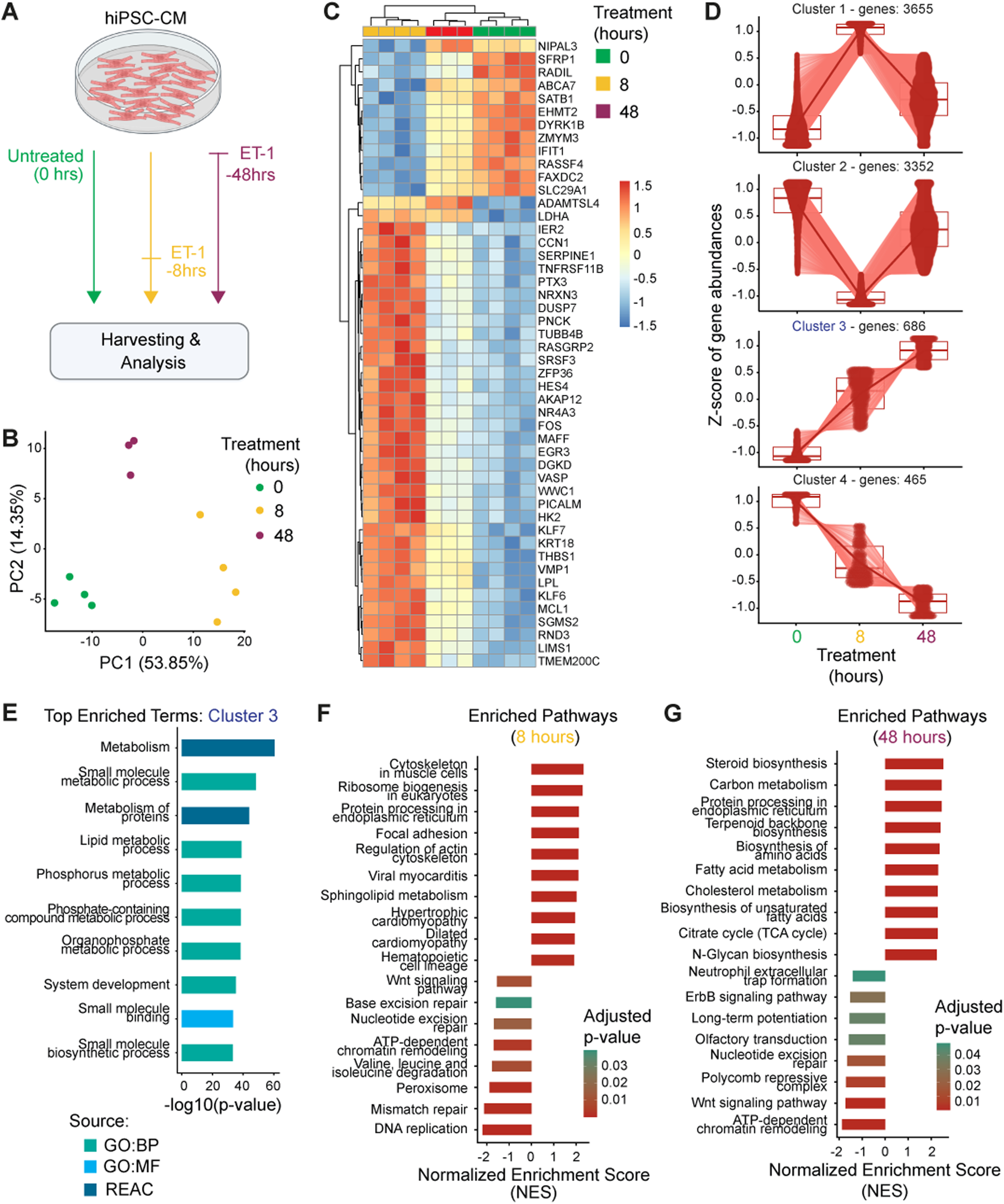
Transcriptomic analysis identifies time-dependent gene expression patterns in hiPSC-CMs treated with ET-1. **A)** Schematic of the experimental protocol for transcriptomic analysis of hiPSC-CMs untreated or treated with ET-1 for 8- or 48-hours. **B)** PCA plot showing the clustering of samples based on duration of ET-1 treatment. **C)** Unsupervised hierarchical clustering of z-scored rlog-transformed gene expression from the top 50 most significantly time-dependent genes reveals strong differences in gene expression profile at each time point. **D)** Clustering analysis of significantly time-dependent genes identifies four distinct patterns of gene expression, using degPatterns from the DEGreport package[50]. For each cluster, the boxplot summarises the gene expression at each time-point, with individual gene-expression overlaid as a density scatter plot. The bold line represents the mean expression trajectory of the cluster. **E)** Top 10 over-represented terms as determined by an over-representation analysis using clusterProfiler of the genes within cluster 3 identify metabolism as a strongly enriched term. Bar length corresponds to the −log_10(_adjusted p value) **F-G)** Selected positively and negatively enriched KEGG pathways as identified with gene-set enrichment analysis using clusterProfiler of genes at 8 (F) and 48 (G) hours reveal a shift in the enriched pathways at each time-point. Bar length represents the normalised enrichment score (NES), and bar colour corresponds to the Benjamini-Hochberg adjusted *P* value.

Gene set enrichment analysis was performed against KEGG pathways on genes ranked by Wald statistic at each treatment timepoint. This identified a transcriptomic programme that changes across the treatment time course. Hour 8 showed upregulation of structural terms (cytoskeleton in muscle cells, regulation of actin cytoskeleton), protein synthesis (protein processing in endoplasmic reticulum, ribosome biogenesis in eukaryotes), and cardiac disease associated terms (hypertrophic cardiomyopathy, dilated cardiomyopathy) (Figure 1F). In contrast, hiPSC-CMs treated with ET-1 for 48 hours were enriched in pathways associated with metabolism (carbon metabolism, biosynthesis of amino acids, cholesterol metabolism) in addition to structural and biosynthetic pathways (Figure 1G).

### 3.2 In-silico modelling of iPSC-CMs treated with ET-1 predicts increased glycolytic and TCA cycle flux, and decreased β-oxidation

To predict changes to the metabolic profile of hiPSC-CMs treated with ET-1, an integrative systems biology approach using both transcriptomic and exometabolomic flux data as input for constraint-based modelling was used. Sample-specific metabolic networks were generated through integration of transcriptomic data with their respective sample-specific media-constrained models. Overall, the reaction availability of these models was similar between groups (Suppl. Figure 4A), consistent with the expectation all three groups will retain largely conserved metabolic networks.

To understand ET-1 associated changes to flux distribution within these metabolic networks, Flux Balance Analysis (FBA) was performed on the sample-specific models. The sample-specific reaction activity identified a broad difference in the metabolic profile of hiPSC-CMs following ET-1 treatment. This identified an increase in glycolysis/gluconeogenesis, particularly at the 8-hour timepoint. A progressive increase in oxidative phosphorylation was also noted in ET-1 treated hiPSC-CMs both at 8 and 48 hours. In contrast to this, a predicted decrease in β-oxidation of even chain fatty acids was observed in ET-1 treated hiPSC-CMs at both 8- and 48-hour time-points. Biosynthetic pathways were also impacted, with an increase in purine metabolism, and decreased pentose phosphate pathway flux (Figure 2A). To examine the central carbon fluxes in more detail a network graph was created showing metabolites as nodes and connecting reactions as edges (Figure 2B). The increased flux through glycolysis with ET-1 treatment appears to be predominantly due to an increase of flux through the hexokinase-1 mediated conversion of glucose to glucose-6-phosphate, the irreversible entry step to glycolysis. Whilst the overall pathway activity score of the TCA cycle decreases with ET-1 treatment, examination of the network model identified an increase in flux through the TCA cycle at 48 hours. The network graph generated for β-oxidation showed a progressive flux decrease over time (Suppl. Figure 4B-D).

**Figure 2.**
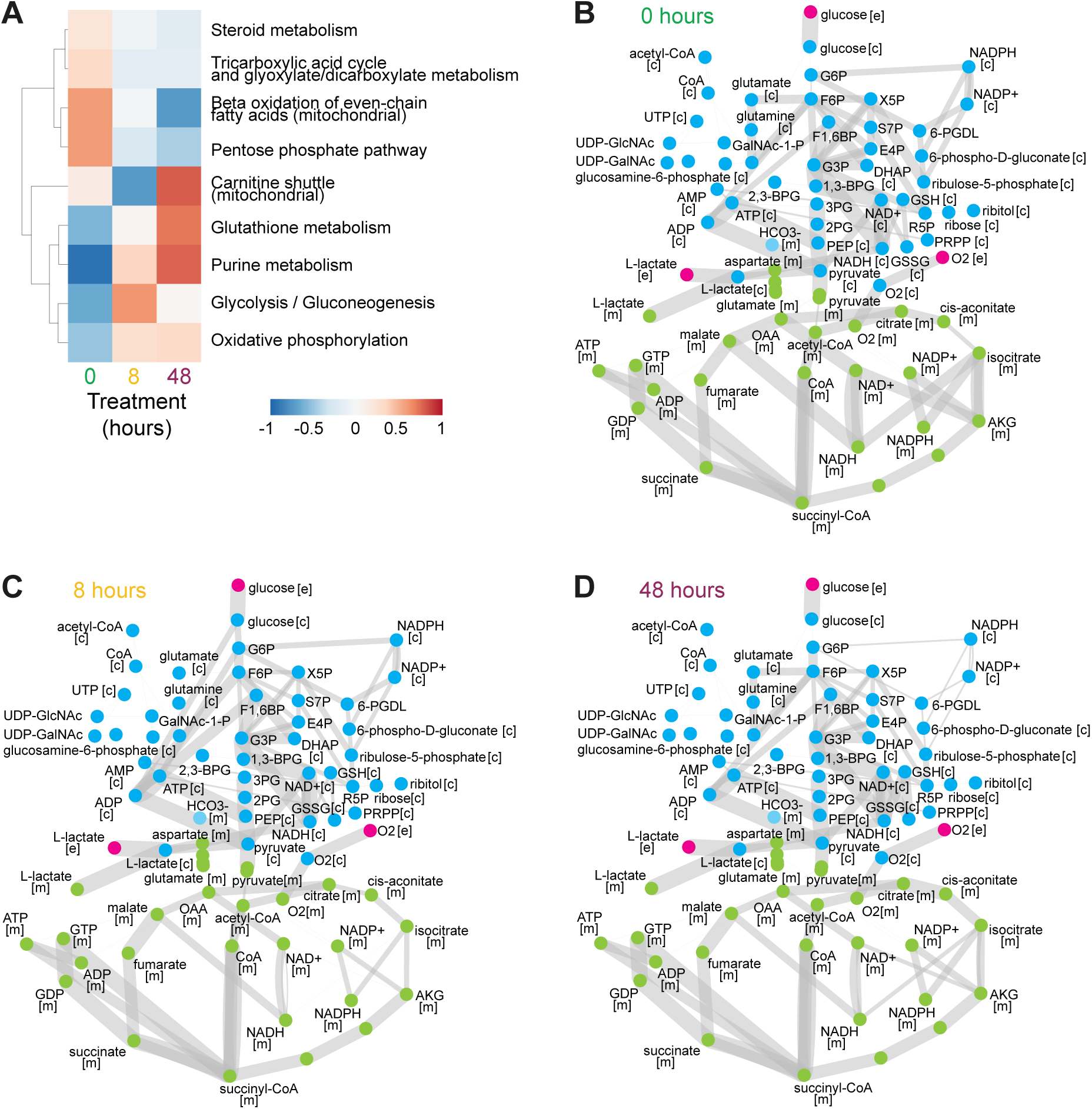
*In silico* modelling predicts changes to central carbon metabolism with ET-1. **A)** Heatmap of pathway activity scores for select pathways between groups, as z-scores. **B-D)** Graphical representations of the predicted fluxes of the metabolic network covering glycolysis, pentose phosphate pathway, hexosamine biosynthesis pathway, and TCA cycle at 0 hours ET-1 (B), 8 hours ET-1 (C), and 48 hours ET-1 (D) using Cytoscape. Nodes represent metabolites, whilst edges represent reactions. Edge width corresponds to the square-root-transformed group mean predicted reaction flux, as described in the Methods.

### 3.3 Extracellular flux analysis confirms model-predicted metabolic rewiring in ET-1-treated hiPSC-CMs

We selected 48 hours of ET-1 treatment as the timepoint for subsequent experiments. At this timepoint, transcriptomic analysis revealed enrichment of metabolic pathways as described above (Figure 1D–E), and hiPSC-CMs also showed clear hypertrophic induction, including increased mRNA expression of *ANP*, *BNP*, *ACTA1* and *MYH6* (Suppl. Figure 5A-D), increased cell surface area (Suppl. Figure 5E), and increased cell volume (Suppl. Figure 5F-G) compared to untreated controls. To experimentally test the metabolic changes predicted by our constraint-based modelling at this timepoint, we next performed extracellular flux analysis using Seahorse instrument. Consistent with the in-silico modelling, the quantification of extracellular acidification rate (ECAR) showed that hypertrophic hiPSC-CMs exhibited significantly elevated glycolytic capacity (Figure 3A-D). The mitochondrial stress test revealed an overall increase in mitochondrial respiration in ET-1-treated hiPSC-CMs compared to untreated controls, including increased basal oxygen consumption rate (OCR), maximal mitochondrial respiration, and ATP-linked OCR (Figure 3E-H). In line with these findings, the OCR over ECAR plot showed that ET-1-treated hiPSC-CMs shifted towards a more metabolically active phenotype compared to untreated cells (Figure 3I). To determine which substrates preferentially drive the increased mitochondrial respiration observed in hypertrophic hiPSC-CMs, we next performed a Seahorse XF substrate oxidation stress test using pathway-specific inhibitors (etomoxir for long-chain fatty acids, UK5099 for glucose/pyruvate, and BPTES for glutamine). Our results showed that the increase in mitochondrial respiration observed is primarily driven by increased glucose oxidation (Figure 3J-K and Suppl. Figure 6).

**Figure 3.**
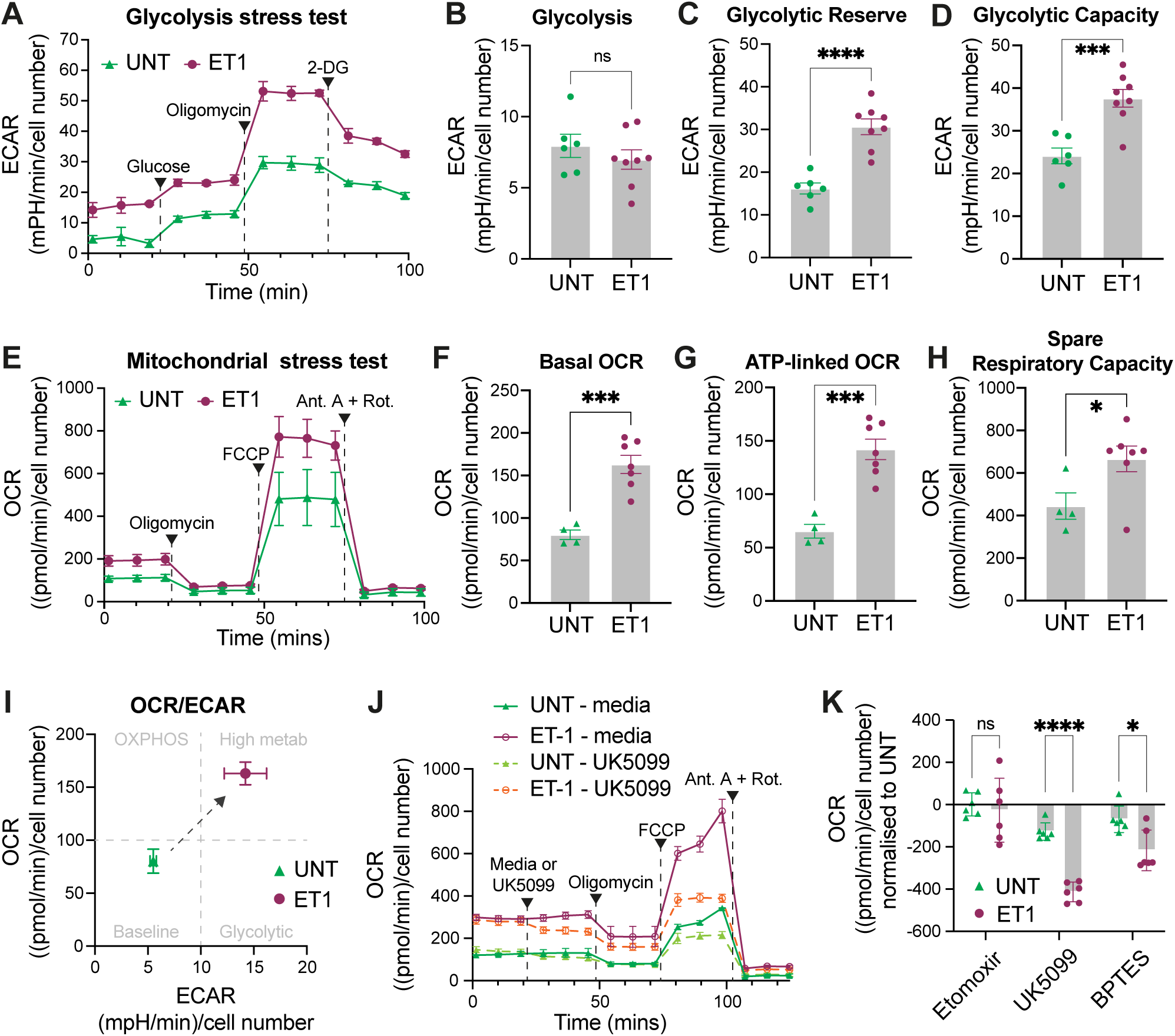
Glycolytic capacity and mitochondrial respiration are enhanced in hypertrophic hiPSC-CMs due to increased glucose utilisation. **A)** Average profile of glycolysis stress test performed in untreated vs. ET-1 treated hiPSC-CMs. **B-D)** quantification of glycolysis, glycolytic reserve, and glycolytic capacity. **E)** Average profiles of mitochondrial stress test measured in untreated vs. ET-1 treated hiPSC-CMs. **F-H)** Quantification of mitochondrial stress test parameters: basal OCR (F), ATP-linked OCR (G) and spare respiratory capacity (H). **I)** OCR over ECAR plot calculated from E). **J)** Average profile of substrate oxidation stress test performed in untreated vs. ET-1 treated hiPSC-CMs either in media or in the presence of the mitochondrial pyruvate carrier inhibitor UK5099. **K)** Quantification of maximal OCR calculated for the substrate oxidation stress in the presence of etomoxir, UK5099 or BPTES. n=3 biological replicates and 2-3 technical replicates/group. Data are presented as mean ± SEM. *P < 0.05, ***P < 0.001, ****P < 0.0001 and ns, not significant by unpaired Student’s t-test in B, C, D and F, G H, and 2-way ANOVA followed by Šidák’s multiple comparison test in K. OCR: oxygen consumption rate; ECAR: extracellular acidification rate.

### 3.4 Stable isotope-resolved metabolomics reveals glucose metabolic remodelling in ET-1-treated hiPSC-CMs

To further validate the observed changes in glucose metabolism, we undertook a SIRM approach using uniformly labelled glucose ([U-^13^C] glucose) followed by targeted LC–MS metabolomics and ^13^C-isotopologue analysis (Suppl. Figure 7A). Targeted intracellular metabolomic analysis via LC-MS on the polar phase identified distinct metabolite abundance profiles for ET-1-treated vs. untreated hiPSC-CMs, as shown by principal component analysis (PCA) (Suppl. Figure 7B). This difference appears to be driven predominantly by changes in amino acids abundance (Suppl. Figure 7C-D), which are required for the protein synthesis to support hypertrophic growth[13, 51], and is consistent with the transcriptomic upregulation of amino acids biosynthesis, whereas overall no clear differences were observed in the abundances of glycolytic or TCA cycle metabolites (Suppl. Figure 7E-F).

SIRM with ^13^C_6-_glucose indicated 48 hours ET-1 treatment results in increased glucose-derived carbon incorporation into several metabolites. Both fractional labelling of the m+6 and m+3 isotopologues and total labelling of glycolytic intermediates were increased in ET-1 treated hiPSC-CMs (Figure 4A-C, Suppl. Figure 8). When we examined incorporation of glucose-derived carbon into intermediates of the glycolytic branch pathways, our results showed that in hypertrophic hiPSC-CMs, there is an increased incorporation of ^13^C in the m+5 isotopologue of pentose phosphate pathway intermediates ribulose-5-phosphate, ribose-5-phosphate and xylulose-5-phosphate (Figure 4D-E and Suppl. Figure 8) and an increased incorporation into the m+3 isotopologue of serine and the m+2 isotopologue of glycine (Figure 4F-G), which make up part of the serine biosynthesis pathway. Glycolysis feeds directly into the tricarboxylic acid (TCA) cycle via pyruvate, and we observed increased m+2 labelling of TCA intermediates including citrate, cis-aconitate, glutamate as a surrogate marker for a-ketoglutarate, succinate, fumarate and malate suggesting an increase in glucose oxidation in ET-1-treated hiPSC-CMs as compared to untreated controls (Figure 4H-K, Suppl. Figure 8). Incorporation at the m+3 isotopologue of TCA intermediates fumarate, malate and aspartate, as surrogate marker for oxaloacetate, was also increased in ET-1-treated hiPSC-CMs, suggesting increased anaplerotic flux from pyruvate to oxaloacetate, bypassing acetyl-CoA (Suppl. Figure 8). In line with our in-silico modelling, which predicted decreased β-oxidation with ET-1 treatment (Figure 2, Suppl. Figure 4B-D), ¹³C-palmitate tracing showed a trend toward a reduction in m+2 labelling of TCA cycle intermediates following ET-1 treatment (data not shown). These findings are compatible with reduced relative palmitate-derived carbon contribution, but do not by themselves quantify fatty acid oxidation rates.

**Figure 4.**
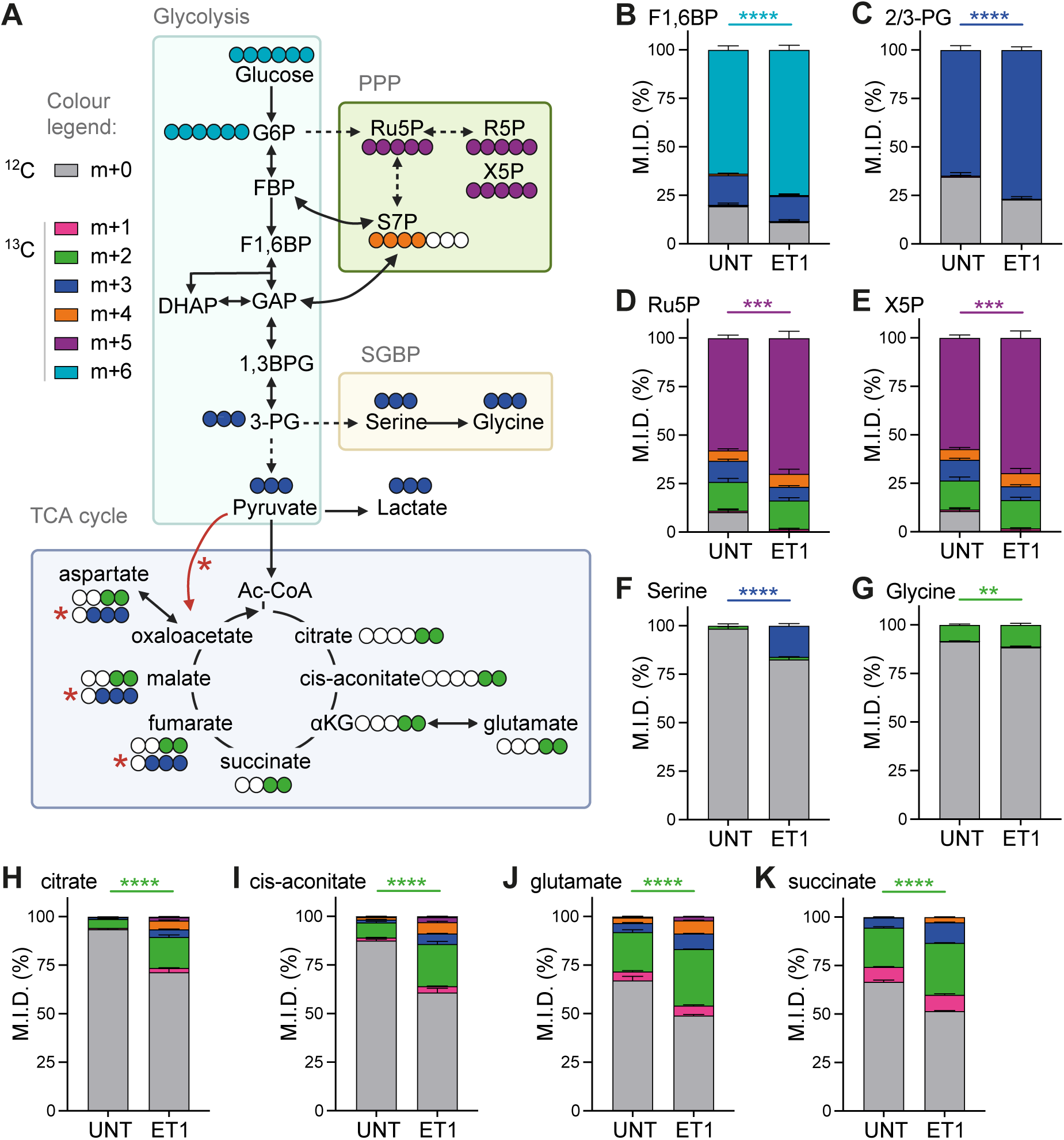
ET-1 treatment induces metabolic reprogramming in hiPSC-CMs, increases glucose flux into glycolysis and anabolic pathways of glycolysis and into the TCA cycle. **A)** Schematic representation of [U-^13^C]glucose carbon labelling through glycolysis, pentose phosphate pathway (PPP), serine biosynthesis pathway(SBP) and TCA cycle. ^13^C-glucose incorporation into **B-C)** glycolytic, **D-E)** pentose phosphate pathway, **F-G)** serine biosynthesis pathway and **H-K)** TCA cycle metabolites in hiPSC-CMs untreated or treated for 48 hrs with ET-1 and for 16 hrs with^13^C-glucose; n=4/group. Data are presented as mean ± SEM. *P < 0.05, **P < 0.01, ***P < 0.001, ****P < 0.0001 by 2-way ANOVA followed by Šidák’s multiple comparison test for mass isotopologue comparisons. M.I.D.: mass isotopologue distribution.

### 3.5 ET-1 induces hypertrophic responses and recapitulates glucose metabolic remodelling in engineered heart tissues

To test whether we can recapitulate hypertrophic and metabolic remodelling in a relevant 3D model, we generated EHTs using a previously described protocol[36]. First, we tested whether ET-1 induces hypertrophic responses in EHTs after 7 days of treatment (Figure 5A). ET-1-treated EHTs showed increased expression of the hypertrophic markers ANP, BNP, ACTA1, and COL1A1 (Figure 5B-E). Chronic ET-1 treatment reduced force amplitude (Figure 5F) and blunted the inotropic responses to isoproterenol challenge relative to untreated controls (Figure 5G-H), as previously described[36]. Having confirmed that ET-1 induces a hypertrophic phenotype in EHTs, we next asked whether the metabolic changes observed in 2D hiPSC-CMs were recapitulated in the 3D EHT platform. Principle component analysis of polar metabolite abundances by targeted LC–MS metabolomics revealed a distinct metabolic profile between the two conditions (Figure 6A). Our results showed an overall increased pool size of metabolites from glycolysis, glycolytic branch pathways, and the TCA cycle in EHTs treated with ET-1 compared to untreated controls (Figure 6B-D). In addition, SIRM using ¹³C-glucose revealed an increased ^13^C labelling of TCA cycle intermediates from glucose, along with increased pyruvate anaplerosis, mirroring the metabolic phenotype observed in 2D hiPSC-CMs (Figure 6E).

**Figure 5.**
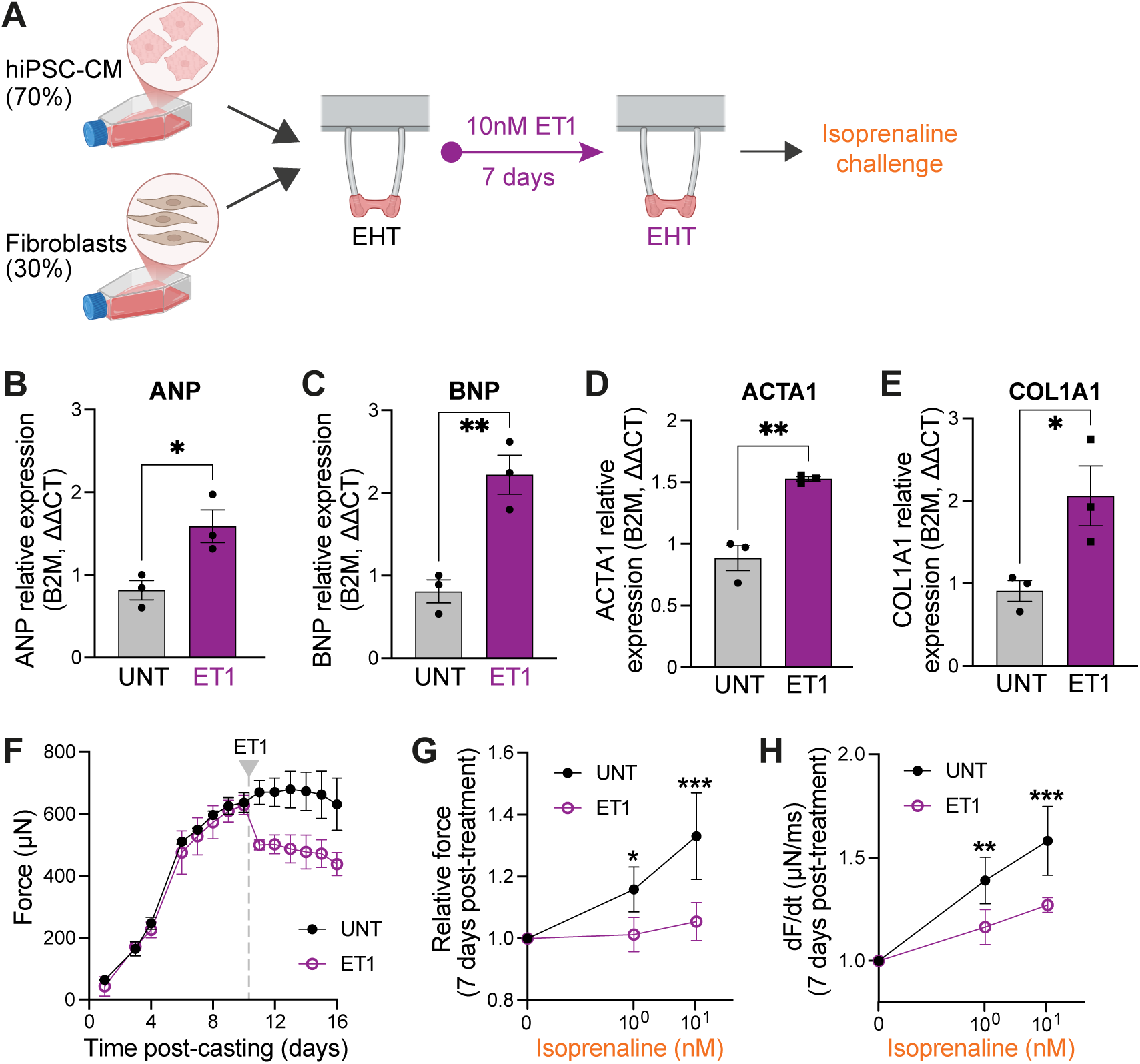
Endothelin-1 treatment induces hypertrophic responses in EHT. **A**) Schematic representation of EHT casting protocol and treatment with 10nM ET-1 for 7 days. **B-E)** Expression levels of ANP, BNP, Acta1, and Col1a1 in ET-1 untreated or treated for 7 days with ET-1. **F)** Force of contraction measured over time. **G)** Relative inotropic response to acute isoproterenol stimulation under 1.5-Hz electrical pacing in CTRL or ET-1-treated EHTs. **H)** Rate of change of force (dF/dt) measured as in G. n=3/group. Data are presented as mean ± SEM. *P < 0.05, **P < 0.01, ***P < 0.001 by unpaired Student’s t-test in B, and 2-way ANOVA followed by Šidák’s multiple comparison test in G and H.

**Figure 6.**
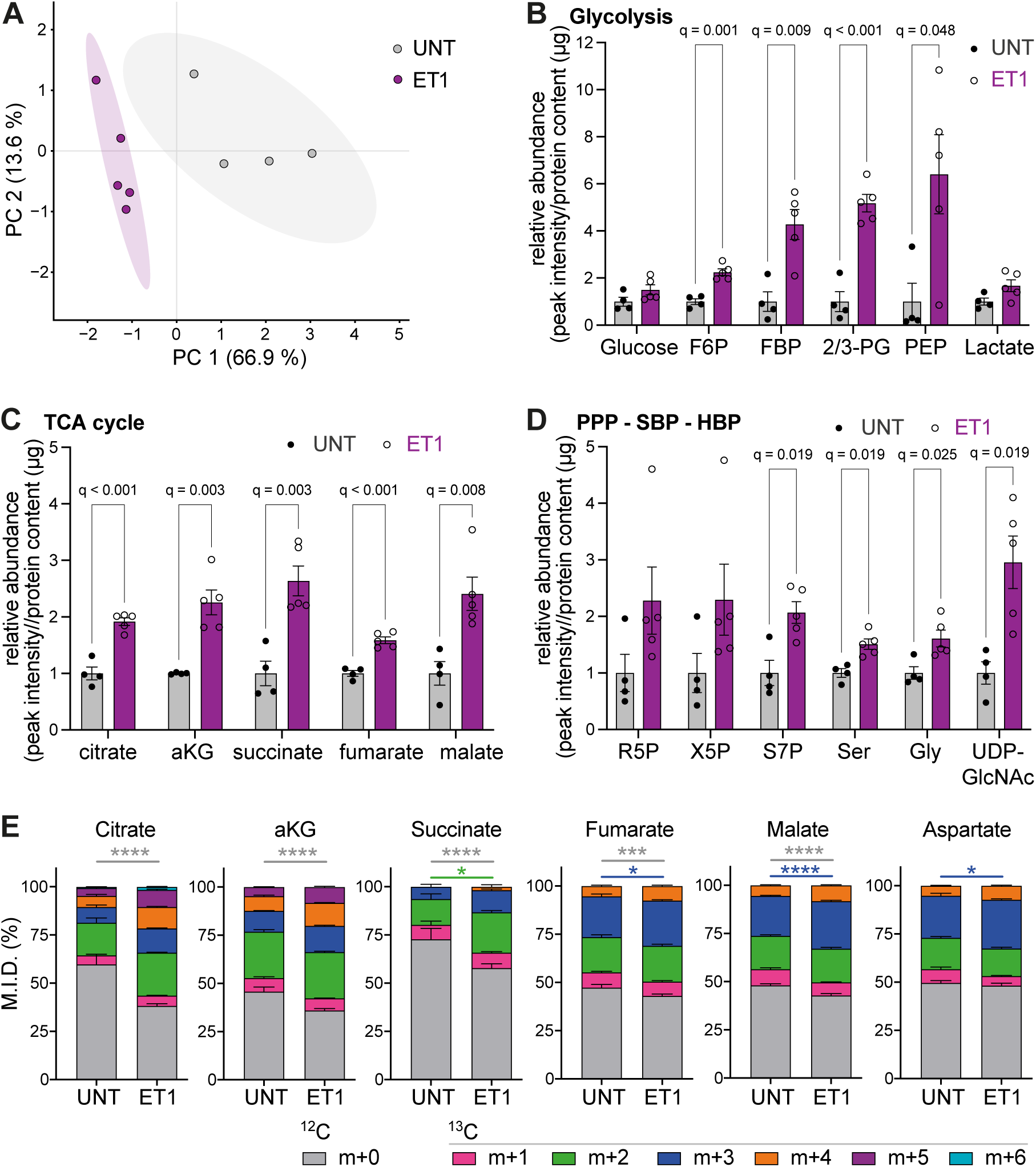
ET1 changes EHT metabolomic profile and increases glucose carbon incorporation into TCA cycle metabolites. **A)** PCA of targeted metabolomic profile of EHTs untreated or ET-1-treated for 7 days. Analysis of glycolytic **B),** TCA cycle **C)** and glycolytic branch pathway **D)** metabolites pool size. **E)** ^13^C-glucose incorporation into TCA cycle metabolites and aspartate as surrogate metabolite for oxaloacetate. n=4-5/group. Data are presented as mean ± SEM. Multiple unpaired t-test with Benjamini-Hochberg correction was performed in B, C and D and 2-way ANOVA followed by Šidák’s multiple comparison test for mass isotopologue comparisons in E. *P < 0.05, **P < 0.01, ***P < 0.001, ****P < 0.0001. TCA: tricarboxylic acid cycle; PPP: pentose phosphate pathway; SBP: serine biosynthesis pathway; HBP: hexosamine biosynthesis pathway; F6P: fructose-6-phosphate; FBP: fructose-1,6-biphosphate; 2/3-PG: 2/3-phosphoglycerate; PEP: phosphoenopyruvate aKG: alpha-ketoglutarate; R5P: ribose-5-phosphate; X5P: xylulose-5-phosphate; S7P: sedoheptulose-7-phosphate; Ser: serine; Gly: glycine; UDP-GlcNAc: uridine-diphosphate N-acetylglucosamine; M.I.D.: mass isotopologue distribution.

## 4.0 Discussion

hiPSC-CMs have proved to be useful models to study human cardiomyocytes in health and disease, and stimulation with ET-1 has previously been shown to recapitulate features of cardiac hypertrophy[23–25, 52]. Metabolic remodelling is recognised as a key part of the chronic response to hypertrophic stimuli such as pressure overload or neurohumoral agonists[53], but there is limited information available regarding metabolic characterisation of hiPSC-CMs treated with ET-1 and the early metabolic response to hypertrophy in a human model. Prior studies have relied on transcriptomic expression of metabolic pathways[23] or lactate secretion as a proxy for anaerobic glycolytic flux[24] which do not provide information on changes in substrate utilisation or contribution of biosynthetic pathways.

Here, using an integrated transcriptomic, metabolomic, stable isotope-resolved metabolomic (SIRM) and extracellular flux approach, we show that ET-1-stimulated hiPSC-CMs in both 2D and 3D culture recapitulate the transcriptional, functional and metabolic responses to hypertrophic induction, including shifts in glucose utilisation that are a key feature in human heart failure.

Our data showed that the transcriptomic response to ET-1 was greatest at the earlier timepoint of 8 hours followed by a decrease at the 48-hour timepoint, consistent with previous reports in both animal cell models[54–56] and hiPSC-CMs[23–25]. Pathway enrichment analysis at each time point recapitulated previously reported responses, with an early enrichment in structural terms involved in hypertrophic response including protein processing and cytoskeleton[25], and later responses predominantly showing enrichment in metabolic pathways[23]. Thus, our model exhibits the expected early transcriptomic response to hypertrophy induction. The transcriptomic response also showed a clear temporal organisation with four distinct expression patterns separating transient changes from those that became progressively more established. Pathway analysis of these clusters showed that genes associated with signalling pathways and cellular component biogenesis increased transiently before returning to baseline (cluster 1), whilst genes associated with metabolic pathways increased progressively and remained elevated (cluster 3) consistent with previous ET-1 studies[23, 25, 55]. This progressive upregulation of metabolic genes suggests a sustained shift in metabolism rather than a transient stress response.

Regulation of metabolism is multifactorial, responding to changes in gene expression, post-translational modifications, co-factor availability, and allosteric regulation[57–60].

Transcriptomics can be used to infer changes to metabolic pathway regulation via gene expression but is insufficient to conclusively predict changes in flux[61]. While metabolomics and SIRM enable examination of specific metabolites and substrate utilisation, these are not yet able to resolve metabolic fluxes across the whole network. In contrast, genome-scale metabolic models can predict which reactions are active and how flux is distributed[62], and provides an efficient approach to integrate multiple omics levels into an in-silico model. In this study, reaction flux distributions were predicted to satisfy the objective function whilst transcriptomic and exchange constraints were used to constrain the solution space. These predicted fluxes were used to evaluate early changes to metabolic flux in hypertrophic remodelling, and predictions were confirmed by in vitro SIRM and Seahorse analyses at the 48-hour timepoint. In line with previous studies, the combined results of in-silico modelling, LC-MS metabolomics, SIRM, and Seahorse identified a substantial metabolic reprogramming during early hypertrophic remodelling characterised by a metabolic switch in substrate utilisation to a predominantly glucose-dependent profile[2, 4, 5]. This metabolic remodelling was accompanied by broad metabolic activation, as evidenced by increased glycolytic activity and mitochondrial respiration as predicted by in-silico flux analysis, measured by extracellular flux analysis, and corroborated by increased ^13^C glucose carbon incorporation into both glycolytic and TCA cycle metabolites. Studies of human cardiac metabolism have reported an increase in myocardial glucose uptake that is initially matched by an increase in glucose oxidation, consistent with the findings presented here[53]. In contrast, some animal models have demonstrated a mismatch between glucose uptake and oxidation[63].

Concomitantly, in-silico modelling predicted a decrease in fatty acids β-oxidation. This prediction was supported experimentally by a trend towards a decrease in palmitate-derived carbon incorporation into TCA cycle metabolites (data not shown), consistent with previous findings[4]. However, inhibition of fatty acid oxidation during extracellular flux analysis did not alter mitochondrial respiration, indicating that these cells are not prioritising fatty acid utilisation. This is likely driven by the standard culture conditions of high glucose and the absence of fatty acids, which favour glucose over fatty acids for energy production[64, 65]. Nevertheless, this study shows that hiPSC-CMs treated with ET-1 enable study of changes to glucose metabolism at the initiation of hypertrophic remodelling.

In addition to canonical conversion of pyruvate to citrate via acetyl-CoA for oxidation in the TCA cycle, pyruvate can enter the TCA cycle via anaplerotic routes, catalysed by either malic enzyme or pyruvate carboxylase. Compatible with increased pyruvate anaplerosis, we observed an increase in m+3 isotopologues of TCA intermediates in the SIRM data, although further studies are required to confirm this observation. Heart failure in human patients has been associated with an increase in pyruvate anaplerosis[66, 67] which may be initially cardioprotective[68] and may support the increased biosynthetic demand of anabolic pathways for protein synthesis[68]. However, pyruvate anaplerosis reduces the efficiency of glucose oxidation and may contribute to oxidative stress through the consumption of NADPH[69]. Whether either consequence applies to this model remains to be tested.

Besides energy production, glucose can be channelled into biosynthesis through a number of ancillary pathways. Our study showed an increase in abundance and glucose incorporation into pathways of intermediary metabolism including the pentose phosphate pathway and serine biosynthesis pathway. This altered glucose contribution to biosynthetic precursor pools in pathological cardiac hypertrophy has been previously described in adult rat cardiomyocytes[13] and mice[70, 71] and has been associated, in those models, with upregulated nucleotide synthesis, glutathione synthesis, and redox regulation in response to cardiac stress. However, the in-silico modelling predicted a slight decrease in flux through both the oxidative- and non-oxidative PPP at both 8 and 48 hours (Figure 2 A and B). This discrepancy may be attributable to the different measurements (glucose-derived carbon contribution vs. flux) or may reflect differences in carbon partitioning or metabolite pool turnover that are not adequately captured by the in-silico models.

Engineered heart tissues improve the maturation of hiPSC-CMs[36]. They also allow for direct measurement of contractile force, which can be coupled to a wide range of endpoint readouts including RNA-seq, qPCR, histology and immunoblotting. Here we extend this approach, to our knowledge for the first time, to LC-MS-based metabolomics and ¹³C glucose isotope tracing.

Decreased responsiveness to isoprenaline and impaired contractility have been previously shown with ET-1 treatment[72], and represent a recognised response to ET-1 in failing human hearts[73]. Here, evaluation of force of contraction and contraction velocity demonstrated a blunting of the inotropic response to isoprenaline stimulation in ET-1 treated EHTs, consistent with these previous reports[73]. Importantly, our LC-MS-based metabolomics and ¹³C glucose isotope tracing results revealed a metabolic response consistent with that observed in the 2D model, characterised by increased glucose oxidation and suggestive of pyruvate anaplerosis.

These findings demonstrate the utility of EHTs for investigating functional changes associated with hypertrophic remodelling and provide additional support for interpreting the associated metabolic changes.

### 4.1 Limitations

hiPSC-CMs are a valuable model of cardiomyocyte function in health and disease, but they retain an immature phenotype. Compared with adult cardiomyocytes, 2D hiPSC-CMs show foetal-like gene expression, immature sarcomere organisation, an absence of t-tubules, and a predominantly glycolytic metabolism with limited fatty acid oxidation[74, 75]. Maturation strategies such as prolonged culture[27], maturation media[76] and 3D cultures including engineered heart tissues[36] improve but do not fully resolve this aspect. Further studies are therefore required to test how metabolic remodelling may differ in metabolically mature hiPSC-CMs, and to better delineate the contribution of fatty acid oxidation to early hypertrophic remodelling.

This study identified unique metabolic profiles in cardiomyocytes undergoing early hypertrophic remodelling using both constraint-based modelling and targeted LC-MS metabolomics. Both approaches involved deliberate methodological choices that define the scope of our conclusions. When performing constraint-based modelling, selecting an appropriate objective function for complex non-proliferative cells can be difficult. In order to satisfy the likely requirements for cardiomyocytes, maximal ATP synthesis was selected, consistent with the energetic demands of contraction, alongside a non-replicative biomass reaction which encompasses protein, RNA, and lipid synthesis. This approach combined with paired transcriptomic and metabolic integration has been recently used to predict metabolic adaptations to cardiotoxicity in a rat cardiac metabolic model[77]. In our study, directionality-based exchange constraints were derived from our targeted metabolomics data. The targeted approach restricts detection to metabolites within our in-house library, but it was chosen because it optimises sensitivity and specificity for the glucose-metabolism intermediates central to this study. Finally, ^13^C partitioning was used as an indicator of pathway activity rather than absolute flux, which would require metabolic flux analysis. Our findings therefore describe the direction of early metabolic remodelling rather than quantitative flux through these pathways.

### 4.2 Conclusion

Taken together, our findings show that ET-1 stimulation drives early and progressive remodelling of glucose metabolism in hiPSC-CMs, characterised by increased glycolytic activity, glucose oxidation and by increased partitioning of glucose carbon into biosynthetic pathways. These changes emerge well before the chronic substrate shift described in animal models of hypertrophy and in failing human myocardium, and they were captured in both 2D monolayers and 3D EHTs, supporting hiPSC-CMs and EHTs as complementary platforms for studying early metabolic remodelling in pathological cardiac hypertrophy. More broadly, coupling genome-scale metabolic modelling with stable isotope-resolved metabolomics provided an integrated framework in which model-predicted flux changes were tested directly against functional and isotope-tracing measurements in a human-relevant system. We anticipate that this framework will extend beyond ET-1, to dissecting metabolic remodelling in other models of hypertrophy and cardiomyopathy and to screening compounds that act on cardiac metabolism.

#### Authors’ contributions

CKTJ: conceptualization, investigation, formal analysis, software, writing - original draft, review & editing; RAK, KKC: conceptualization, investigation, formal analysis, writing - review & editing; EdS: investigation, writing – review & editing; RRO, EB, EK, SV: investigation; CMR: investigation, formal analysis, methodology; CXCS: methodology; KSB: resources, writing - review & editing; AMS: conceptualization, funding acquisition, supervision, writing - review & editing; AZ: conceptualization, formal analysis, funding acquisition, supervision, writing – original draft, review & editing. All authors approved the final manuscript.

## Acknowledgements

Figure 1A, 4A, 5A, Suppl. Fig 1A, Suppl. Fig 2A, Suppl. Fig. 7A were created using BioRender licensed to Anna Zoccarato.

---

**Suppl. Figure 1.**
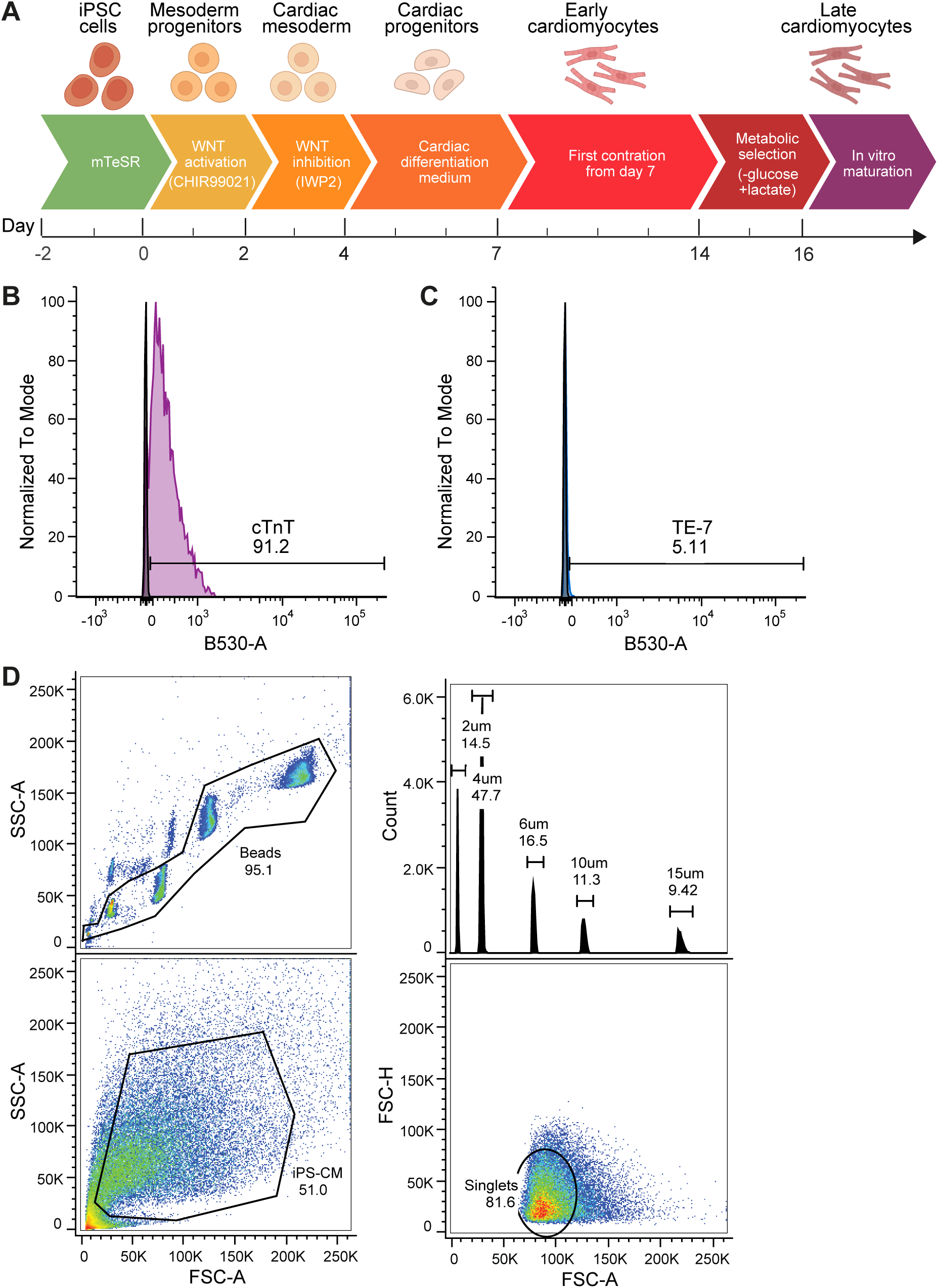
**A)** Schematic of the hiPSC-CM differentiation protocol. **B-C)** FACS analysis of hiPSC-CM at the end of the differentiation protocol showing the percentage of cells positive for cTnT (B) or TE-7 (C). **D)** Top Left: Forward scatter (FSC) vs side scatter (SSC) dot plot showing five populations of size-calibration beads, debris and clumps were excluded by gating. Top right: FSC area (FSC-A) histogram used to plot calibration curve. Bottom left: FSC vs SSC dot plot showing hiPSC-CMs population gate (black) Bottom right: FSC height (FSC-H) vs FSC-A histogram with black gate used for doublets exclusion.

**Suppl. Figure 2.**
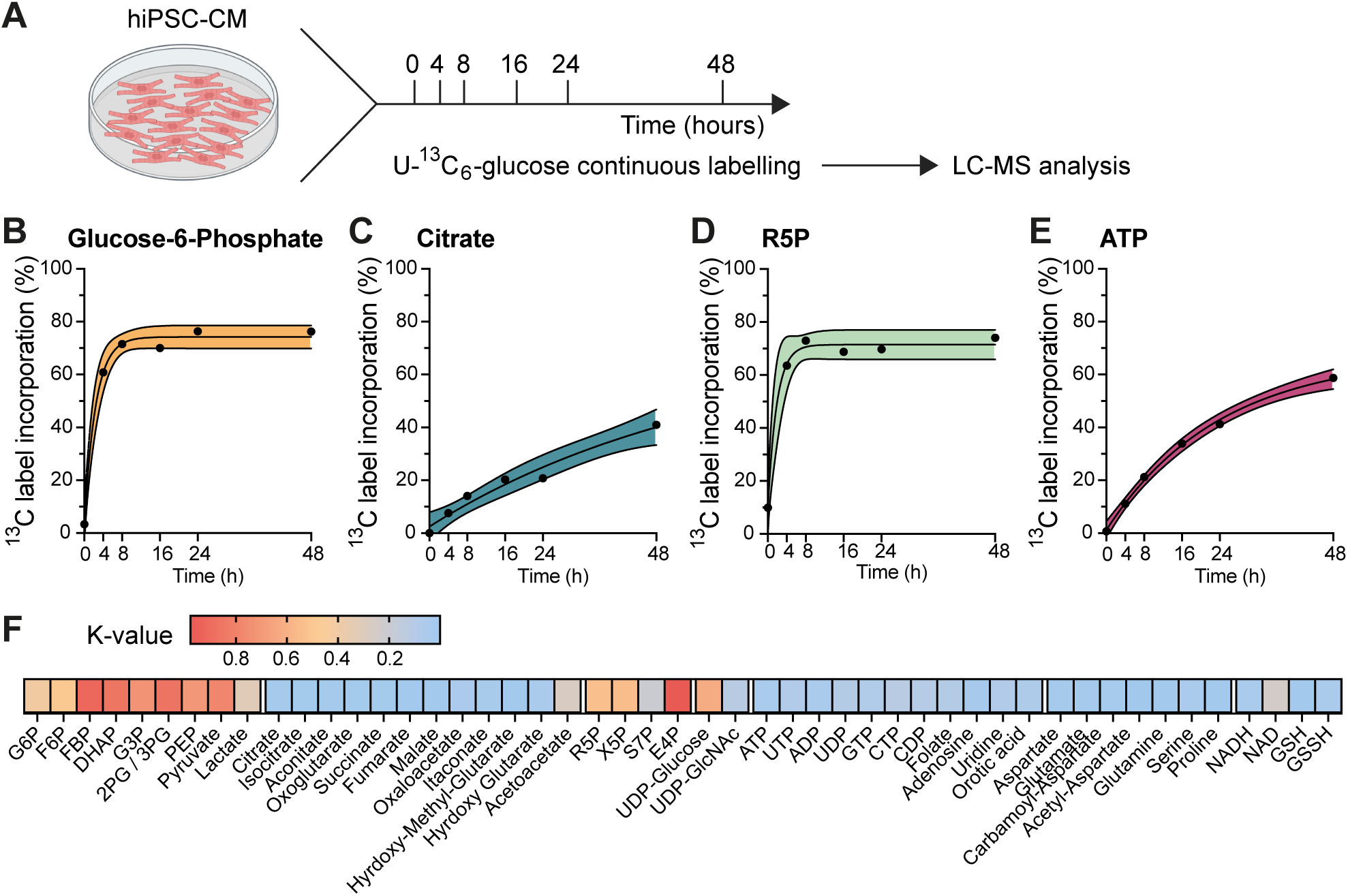
**A)** Outline of the experimental workflow for the time course analysis of ^13^C_6 g_lucose tracing in hiPSC-derived cardiomyocytes. **B)** Representative labelling time courses for metabolites (G6P, citrate, R5P and ATP) exhibiting different label incorporation paces or glucose contribution. (n = 3 biological replicates in each time point); black dot represents the mean ^13^C label incorporation in %. **C)** Heat map displaying average k-values for metabolites linked to selected pathways of interest. The labelling rate k (per hour) represents the rate constant derived from fitting data to a first-order exponential model in GraphPad Prism.

**Suppl. Figure 3.**
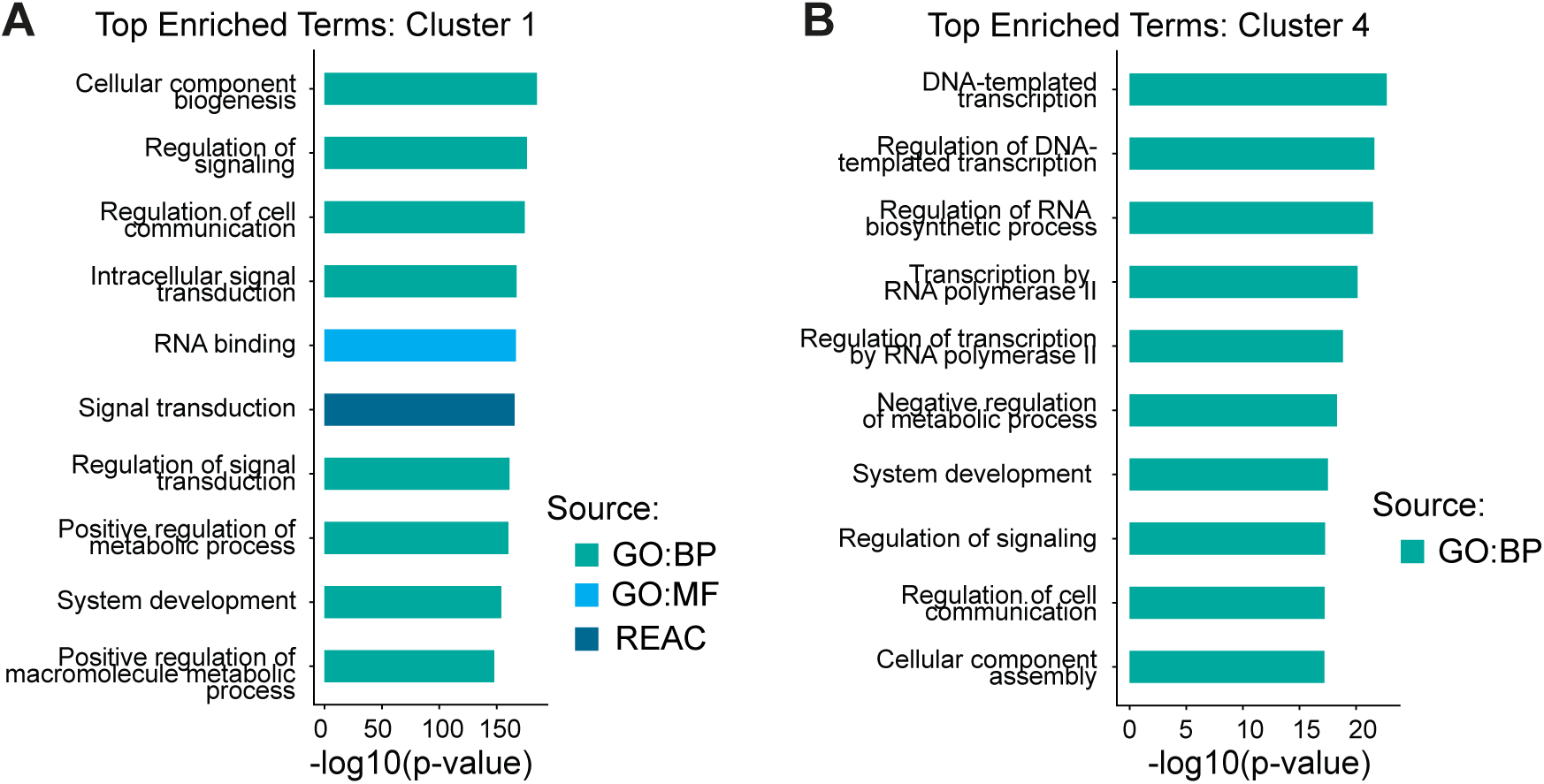
Over-representation analysis of gene clusters based on time-dependent expression reveals functional differences between expression clusters. Bar plot of the top 10 most significantly over-represented terms for cluster 1 (A), and cluster 4 (B) identified by over-representation analysis using clusterProfiler. Bar length indicates –log_10(_adjusted p-value). GO:BP; gene ontology: biological process, GO:MF; gene ontology: molecular function, REAC; Reactome pathway.

**Suppl. Figure 4.**
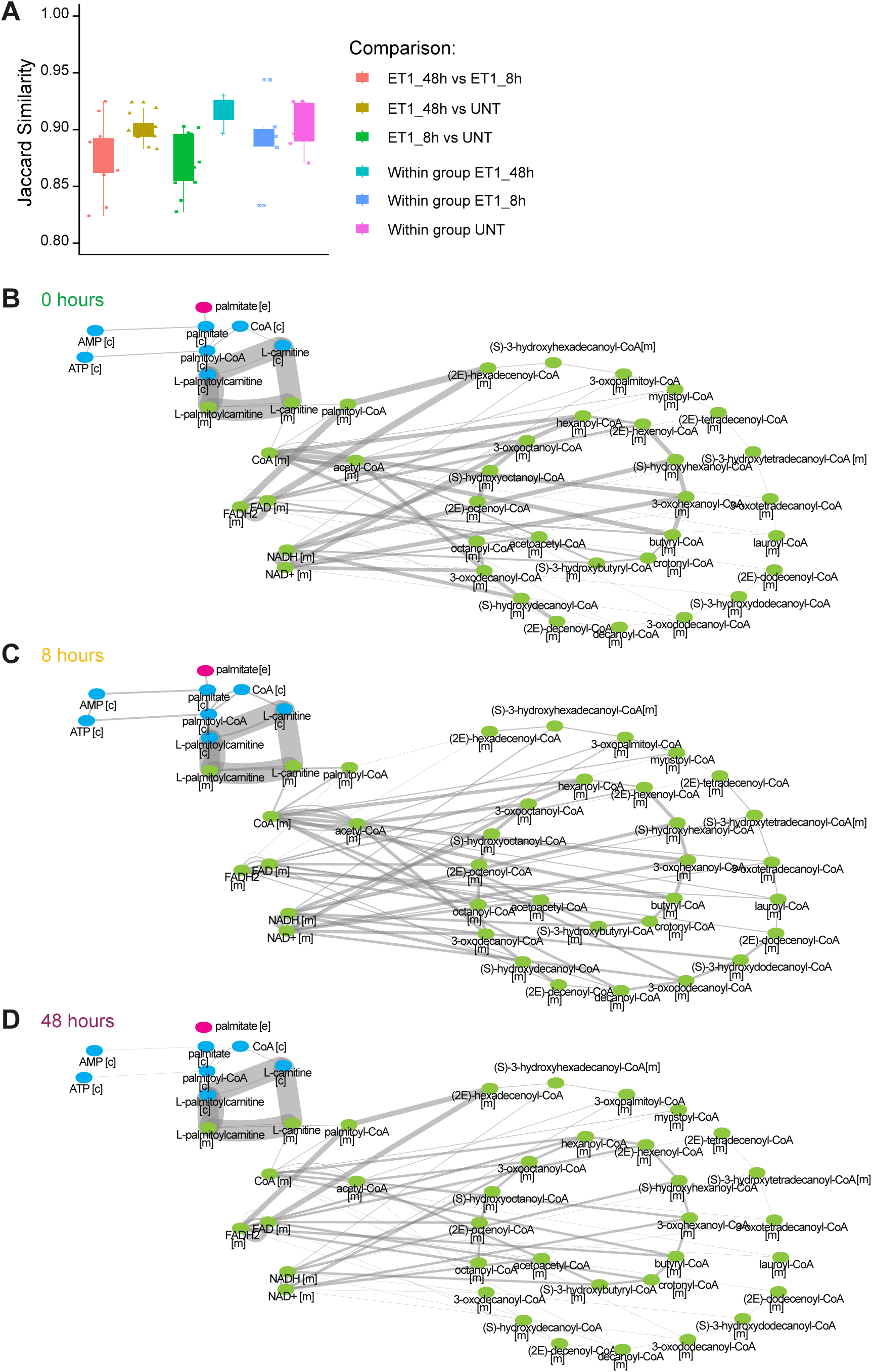
In silico modelling predicts changes to metabolic flux with ET-1 treatment. **A)** Structural variation between models from untreated, 8 hours ET-1, and 48 hours ET-1, as calculated by the Jaccard similarity index. Graphical representation of mitochondrial β-oxidation of even chain fatty acids shown as mean metabolic fluxes in **B)** untreated, **C)** at hour 8 and **D)** at hour 48 of ET-1 treatment. Nodes represent metabolites, whilst edges represent reactions. Edge width corresponds to square-root transformed predicted reaction flux magnitude.

**Suppl. Figure 5.**
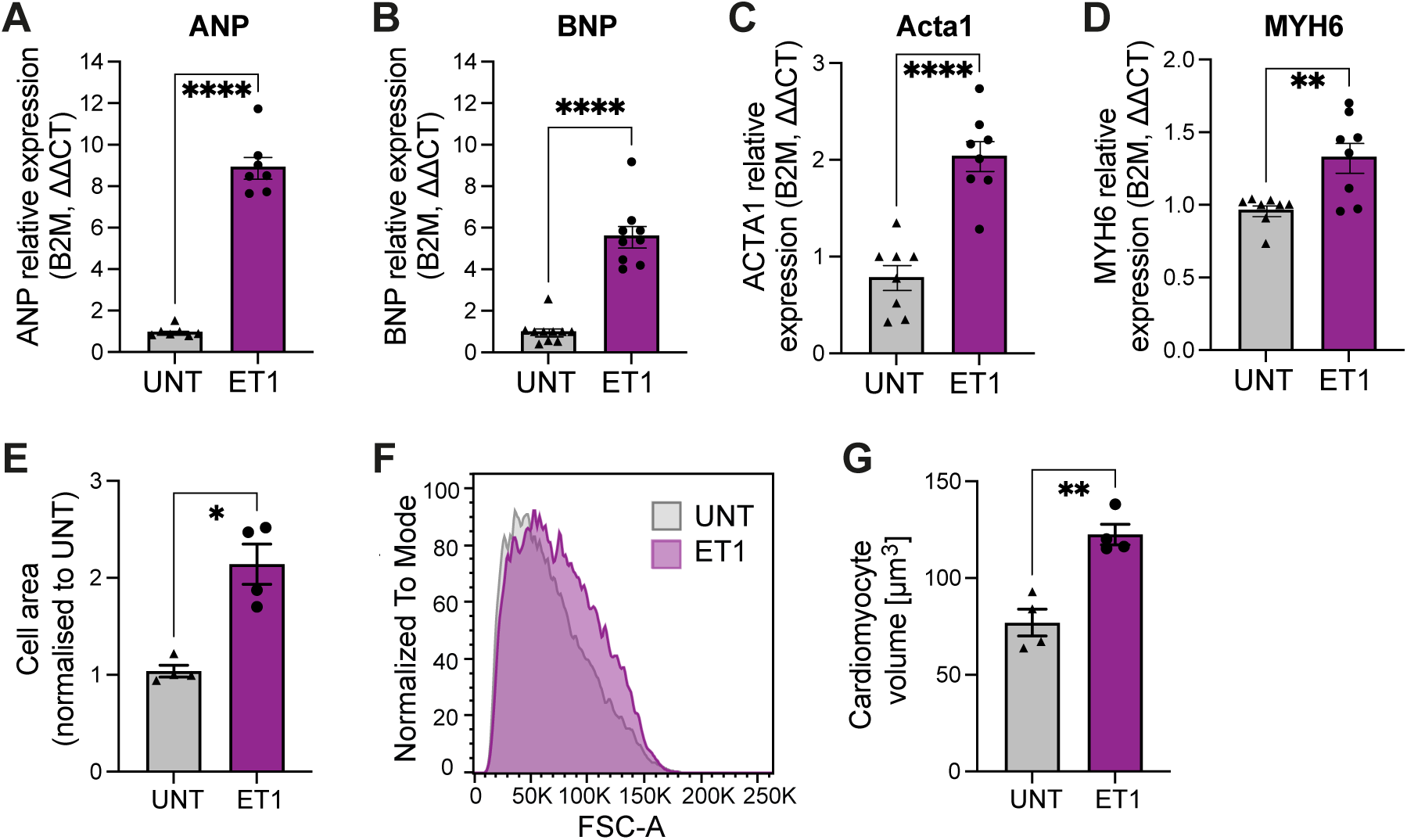
48 hours ET-1 treatment induces hypertrophic responses in hiPSC-CMs. Expression levels of **A)** ANP**, B)** BNP, **C)** ACTA1 and **D)** MYH6 in hiPSC-CMs treated for 48 hours with ET-1; n=7-8/group. **E)** Cell surface area quantification of hiPSC-CMs untreated or ET-1-treated for 48 hours. **F)** FACS analysis of hiPSC-CMs volume as described in Methods. **G)** Quantification of hiPSC-CMs cell volume by FACS. n=4/group. Data are presented as mean ± SEM. *P < 0.05, **P < 0.01, ns, not significant by unpaired t-test.

**Suppl. Figure 6.**
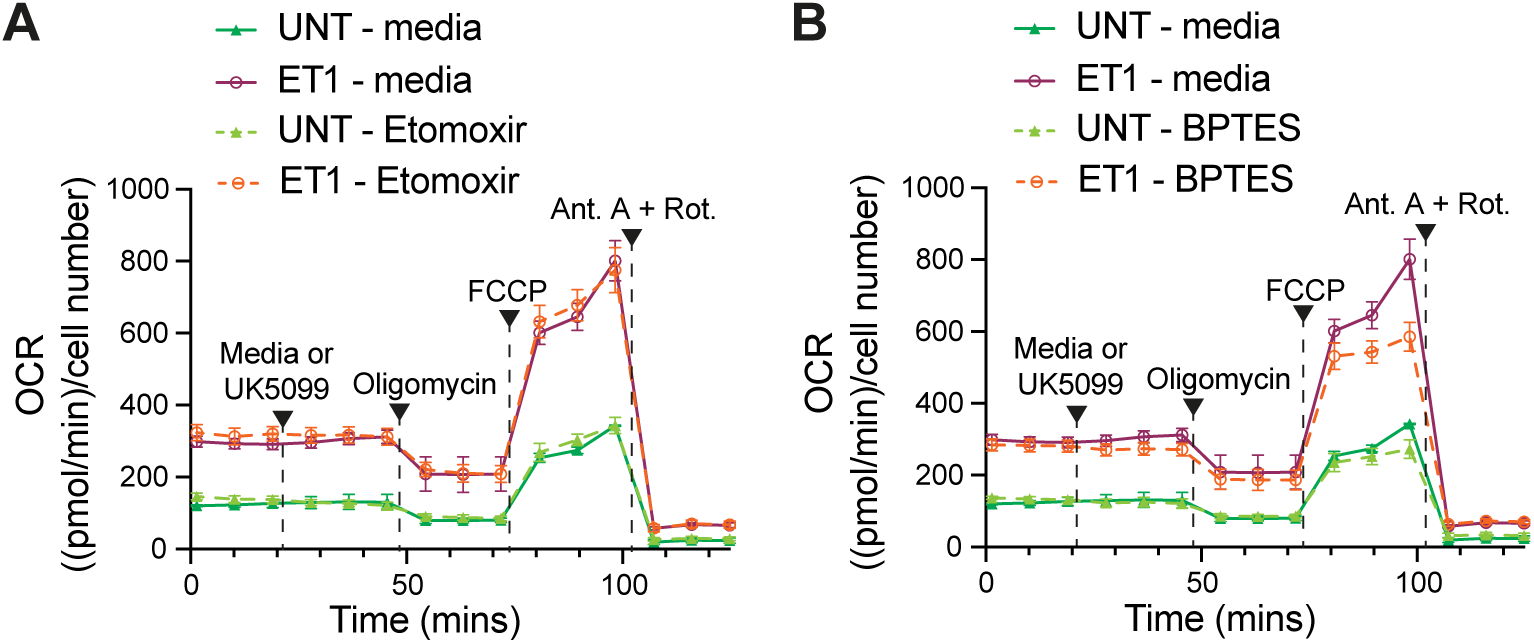
Average profile of substrate oxidation stress test performed in untreated vs. ET-1 treated hiPSC-CMs adding as first injection media or etomoxir **(A)**, or BPTES **(B)**. n=3 biological replicates and 2-3 technical replicates/group.

**Suppl. Figure 7.**
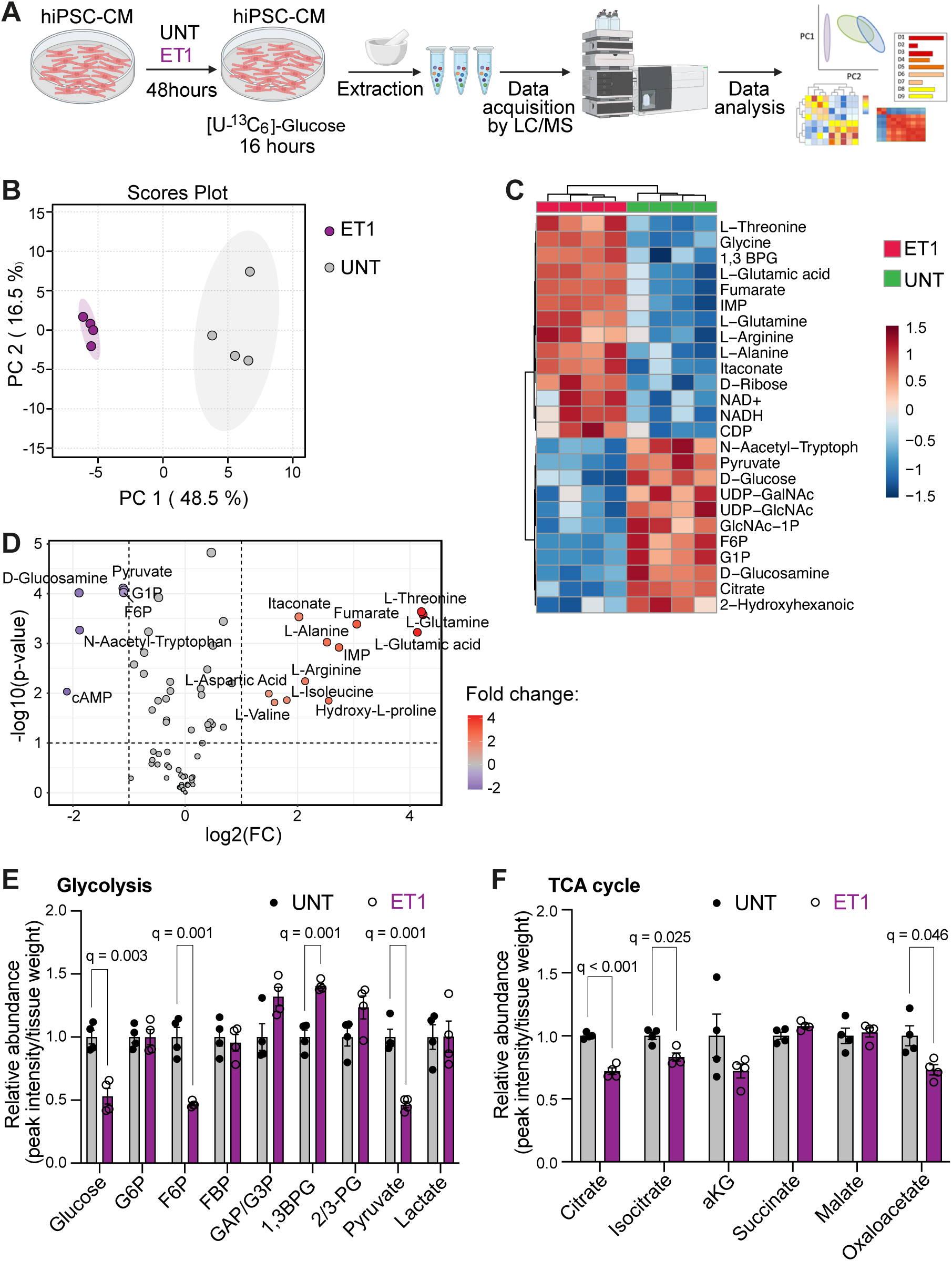
**A**) Schematic representation of hiPSC-CMs hypertrophic induction coupled with ^13^C_6-_glucose incubation followed by targeted LC–MS metabolomics and ^13^C-isotopomer analysis. **B)** Principal component analysis of targeted metabolomic profile of untreated vs. ET-1 treated hiPSC-CMs, n ≥ 4/group. **C)** Heat map showing hierarchical clustering of metabolite abundance for the top 25 features in untreated vs. ET-1 treated hiPSC-CMs. **D)** Important features selected by volcano plot with fold change threshold (x) 2 and t-tests p-value threshold (y) 0.1. Blue: decreased, red: increased, in ET-1 treated hiPSC-CMs. Analysis of glycolytic **E)** and TCA cycle **F)** metabolites abundance. Data are presented as mean ± SEM. q was calculated by multiple unpaired t-test with Benjamini-Hochberg correction in E and F. LC-MS: liquid chromatography/mass spectrometry; 1,3 BPG: 1,3-biphosphoglycerate; IMP: inosine monophosphate; NAD+: nicotinamide adenosine dinucleotide; NADH: reduced nicotinamide adenosine dinucleotide; CDP: cytidine diphosphate; UDP−GalNAc: uridine diphosphate-N-acetylgalactosamine; UDP-GlcNAc: uridine-diphosphate N-acetylglucosamine; GlcNAc-1P: N-acetyl-α-D-glucosamine 1-phosphate; F6P: fructose-6-phosphate; G1P: glucose-1-phosphate; G6P: glucose-6-phosphate; FBP: fructose-1,6-bisphosphate; GAP/G3P: glyceraldehyde-3-phosphate/glycerol-3-phosphate; 2/3-PG: 2/3-phosphoglycerate; aKG: alpha-ketoglutarate.

**Suppl. Figure 8.**
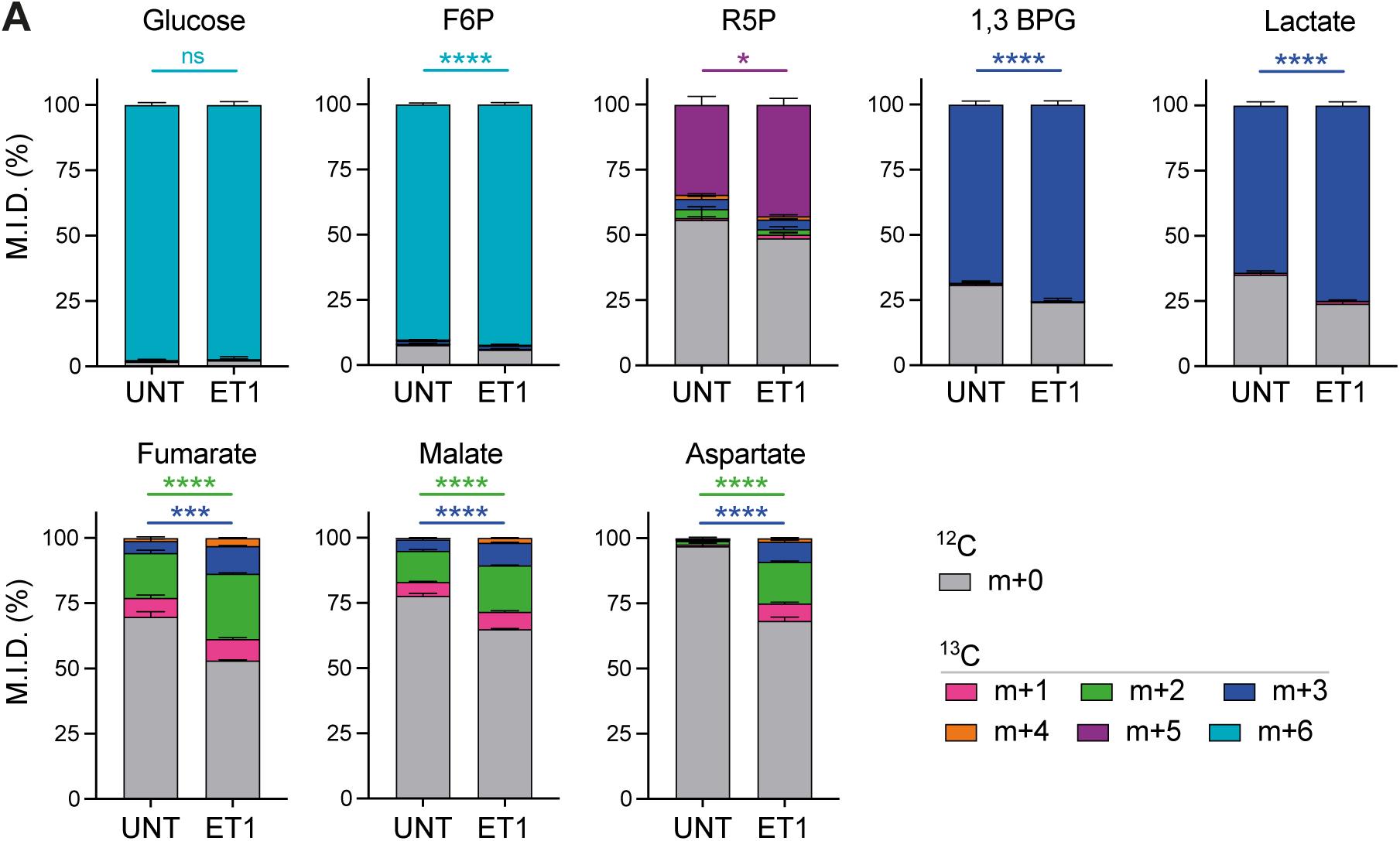
^13^C-glucose incorporation into glycolytic (glucose, F6P, 1,3BPG, lactate), pentose phosphate pathway (R5P) and TCA cycle (fumarate, malate, aspartate as surrogate for oxaloacetate) metabolites in hiPSC-CM untreated or treated for 48 hrs with ET-1 and for 16 hrs with^13^C-glucose; n=4/group. Data are presented as mean ± SEM. *P < 0.05, **P < 0.01, ***P < 0.001, ****P < 0.0001, ns, not significant by 2-way ANOVA followed by Šidák’s multiple comparison test for mass isotopologue comparisons. F6P: fructose-6-phosphate; 1,3BPG: 1,3-bisphosphoglycerate; R5P: ribose-5-phosphate; TCA: tricarboxylic acid. M.I.D.: mass isotopologue distribution.

| RPMI 1640 |  |
|---|---|
|  | L-Arginine |
|  | L-Asparagine |
|  | L-Aspartic acid |
|  | L-Glutamate |
|  | L-Glutamine |
|  | Glycine |
|  | L-Histidine |
|  | L-Isoleucine |
|  | L-Leucine |
|  | L-Lysine |
|  | L-Methionine |
|  | L-Phenylalanine |
|  | L-Proline |
|  | L-Serine |
|  | L-Threonine |
|  | L-Tryptophan |
|  | L-Tyrosine |
|  | L-Valine |
|  | L-Cystine |
|  | Trans-4-hydroxy-L-proline |
|  | Glutathione |
|  | D-Glucose |
|  | Phenol Red |
|  | Chloride |
|  | Calcium |
|  | Potassium |
|  | Sulfate |
|  | Magnesium |
|  | Bicarbonate |
|  | Sodium |
|  | Phosphate |
|  | Calcium Nitrate |
|  | Magnesium Sulfate |
|  | Potassium Chloride |
|  | Sodium Bicarbonate |
|  | Sodium Chloride |
|  | Sodium Phosphate dibasic |
|  | p-Aminobenzoate |
|  | D-Biotin |
|  | Choline |
|  | Folate |
|  | myo-Inositol |
|  | Niacinamide |
|  | D-Calcium pantothenate |
|  | Pyridoxin |
|  | Riboflavin |
|  | Thiamine |
|  | Vitamin B12 |
| B27 |  |
|  | Biotin |
|  | Dl-alpha-tocopherol acetate |
|  | DL-alpha-Tocopherol |
|  | Retinol, acetate |
|  | Bovine Serum Albumin |
|  | Albumin |
|  | Catalase |
|  | Transferrine |
|  | Superoxide Dismutase |
|  | D-Galactose |
|  | Ethanolamine Hydrochloride |
|  | Glutathione |
|  | Levocarnitine Chloride |
|  | Linoleate |
|  | Linolenic Acid |
|  | Putrescine Dihydrochloride |
|  | Sodium Selenite |
|  | Liothyronine |
|  | Progesterone |
|  | Corticosterone |

